# A Pooled Cell Painting CRISPR Screening Platform Enables de novo Inference of Gene Function by Self-supervised Deep Learning

**DOI:** 10.1101/2023.08.13.553051

**Authors:** Srinivasan Sivanandan, Bobby Leitmann, Eric Lubeck, Mohammad Muneeb Sultan, Panagiotis Stanitsas, Navpreet Ranu, Alexis Ewer, Jordan E. Mancuso, Zachary F Phillips, Albert Kim, John W. Bisognano, John Cesarek, Fiorella Ruggiu, David Feldman, Daphne Koller, Eilon Sharon, Ajamete Kaykas, Max R. Salick, Ci Chu

## Abstract

Pooled CRISPR screening has emerged as a powerful method of mapping gene functions thanks to its scalability, affordability, and robustness against well or plate-specific confounders present in array-based screening ^1–6^. Most pooled CRISPR screens assay for low dimensional phenotypes (e.g. fitness, fluorescent markers). Higher-dimensional assays such as perturb-seq are available but costly and only applicable to transcriptomics readouts ^7–11^. Recently, pooled optical screening, which combines pooled CRISPR screening and microscopy-based assays, has been demonstrated in the studies of the NFkB pathway, essential human genes, cytoskeletal organization and antiviral response ^12–15^. While the pooled optical screening methodology is scalable and information-rich, the applications thus far employ hypothesis-specific assays. Here, we enable hypothesis-free reverse genetic screening for generic morphological phenotypes by re-engineering the Cell Painting ^16^ technique to provide compatibility with pooled optical screening. We validated this technique using well-defined morphological genesets (124 genes), compared classical image analysis and self-supervised learning methods using a mechanism-of-action (MoA) library (300 genes), and performed discovery screening with a druggable genome library (1640 genes) ^17^. Across these three experiments we show that the combination of rich morphological data and deep learning allows gene networks to emerge without the need for target-specific biomarkers, leading to better discovery of gene functions.

## Introduction

CRISPR-based genetic screens allow researchers to causally connect genes to their cellular phenotypes and functions. While such studies can be conducted in array format, pooled CRISPR screening methodologies, stemming from seminal work utilizing barcoded shRNA technology ^18,19^, are generally more cost-effective and scalable ^6^. Pooled CRISPR screens typically assay for low dimensional readouts, such as cell survival ^1,2,5,20^ or sortable fluorescent biomarkers ^21–23^. While these CRISPR screens can produce valuable insights, the limited dimensionality of readouts leads to a reliance on a well defined phenotypic marker, which prevents hypothesis-free exploration. Perturb-seq was developed to combine high dimensional single cell RNAseq phenotypes with pooled CRISPR screening ^7–11^; however it is cost prohibitive at large scales. In addition, transcriptomic data do not fully capture cell state and morphological data can provide complementary information in MoA predictions ^24^.

Recently, pooled optical screening methods have employed bespoke phenotypic assays designed for studying specific biological questions such as the NFkB pathway, antiviral response, cytoskeletal organization, and essential genes ^12–15^. In contrast, a generic morphological assay would enable hypothesis-free biological exploration. Inspired by the Cell Painting assay ^16^, which has been successfully applied towards clustering genes by similar functions ^25^, virtual screening for small molecules ^26^ and MoA prediction ^24,27^, we sought to address the current incompatibility between pooled optical screening and Cell Painting, which includes the spectral collision between Cell Painting and 4-color *in situ* sequencing (ISS) and the RNA degradation caused by the Cell Painting workflow. By combining Cell Painting with pooled optical screening, we aim to build a platform that would provide datasets conducive for machine learning (ML) ^28^, enable unbiased mapping of genes to functions, and help improve drug discovery.

We report here Cell Painting Pooled Optical Screening in Human cells (CellPaint-POSH). We modified the original Cell Painting technique in several ways to provide compatibility with ISS. We compared self-supervised ML representation learning with classical image analysis, and demonstrated that machine learnt features enable better prediction tasks than expert-engineered morphological features ^29^. We demonstrated that morphological phenotypes successfully cluster genes by known functions, and that this unbiased profiling method is capable of discovering genetic functions and networks and uncovering new insights without explicit pathway-specific reporters.

## Results

### Development and optimization of CellPaint-POSH

We made several modifications to optimize the Cell Painting assay ^16^ for pooled optical screening (workflow described in Fig. 1a, detailed plasmid maps for lentiviral delivered Cas9 and single-guide RNA (sgRNA) shown in Fig. S1a-b). First, MitoTracker used in Cell Painting is a stress-inducing live cell stain, and the sustained fluorescence of MitoTracker into the sequencing stages prohibited reliable *in situ* base calling (data not shown). We thus developed an RNA-based label for mitochondria that we term “Mitoprobe.” Briefly, using the structure of the human mitochondrial ribosome from the Protein Data Bank (3J9M) ^30^, we identified the most likely RNA sequences to bind to the human mitochondrial ribosome’s 12s and 16s ribosomal RNA (rRNA) (Fig. 1b). We then optimized on solvent-accessible surface area (SASA) ^31^, probe length, GC content, absence of repetitive bases, and intra-sequence distance (Fig. S1c), and conjugated the 5’ end with Cy5. Simultaneous co-staining of fixed adenocarcinomic human alveolar basal epithelial (A549) cells with Mitoprobe and Tom20 antibody gives concordant staining patterns (Fig. 1c, Fig. S1d), demonstrating in a POSH-compatible mitochondrial RNA probe that can be applied to fixed cells (Table 1), streamlining all cellular stains into a single step in the workflow. The RNA-FISH based mitochondrial label is washed out during ISS chemistry and heating steps, avoiding optical interference with 4-color sequencing. Second, RNAse inhibitor Ribolock was implemented during all staining phases to prevent degradation of mitochondrial rRNA, sgRNA spacer containing transcripts and other cellular RNAs. Third, we observed that subjecting cells to Cell Painting prior to reverse transcription (RT) led to degradation of RNAs and ISS signals (data not shown). Moving RT prior to Cell Painting preserved RNAs and led to successful ISS (Fig. 1d).

**Figure 1.**
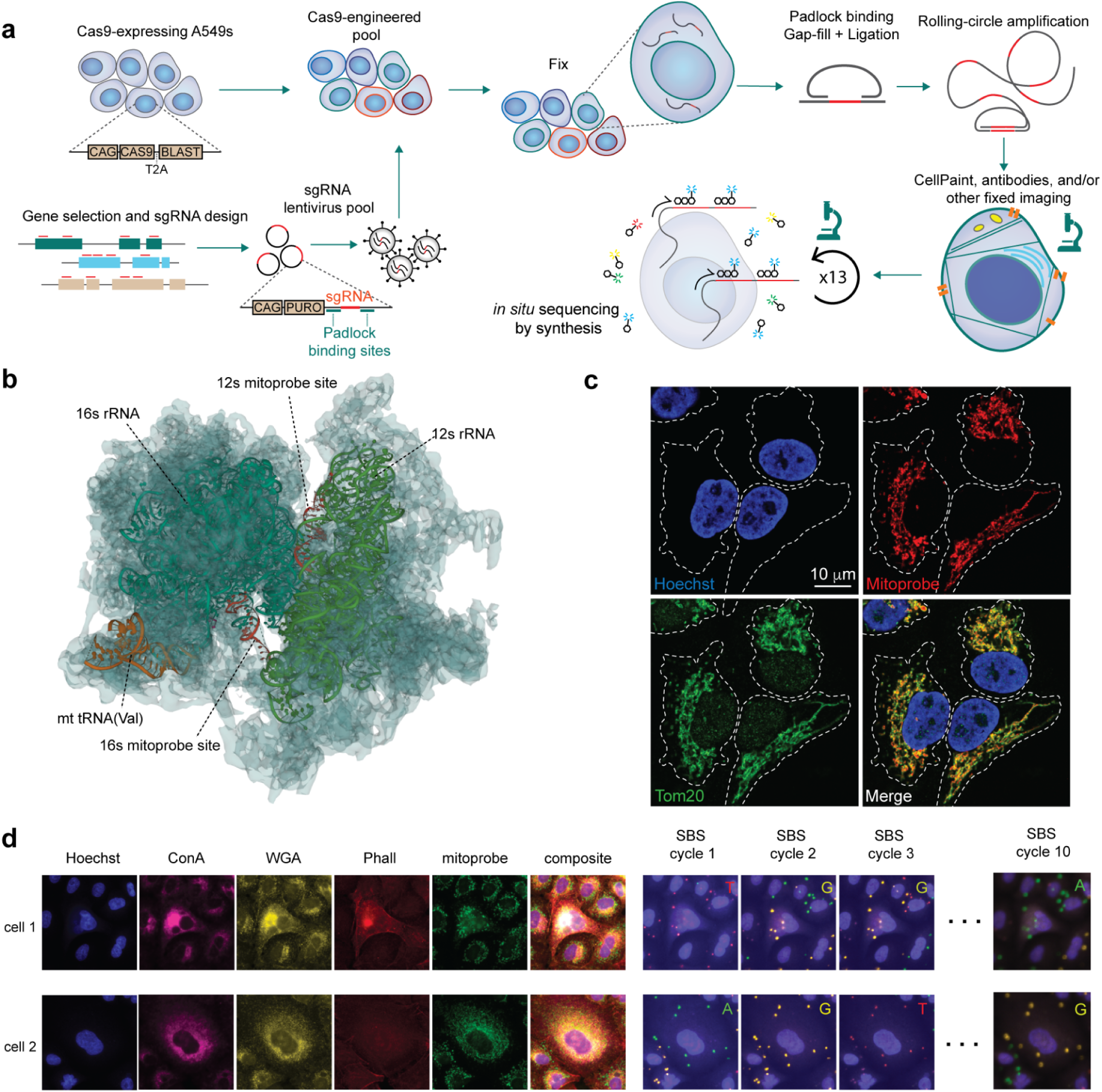
Overview and development of POSH platform. **a**. a general overview of the Pooled Optical Screening in Human cells platform (POSH). A pool of cells is generated that contain both constitutively-active Cas9, as well as lentivirally-delivered sgRNAs targeting a select number of genes. Cells are fixed and the sgRNA sequences are amplified via rolling circle amplification (RCA). The CellPaint imaging assay is then conducted on the cells followed by *in situ* sequencing by synthesis to match image data with transfected sgRNA and gene knockout. **b**. the protein structure of the human mitochondrial ribosome (transparent), 16s, 12s, and Val highlighted in teal, green, and orange, respectively. The exposed regions of the 12s and 16s rRNAs are highlighted in red. **c**. 60X water-immersion confocal fluorescence images of A549s treated with Hoechst, mitoprobe, and anti-Tom20 antibody, reveals close overlap between mitoprobe and Tom20 without nuclear staining. **d**. example raw images showing the multiple CellPaint tiles, as well as the several sequencing-by-synthesis (SBS) tiles paired to the phenotypic data.

**Table 1:**
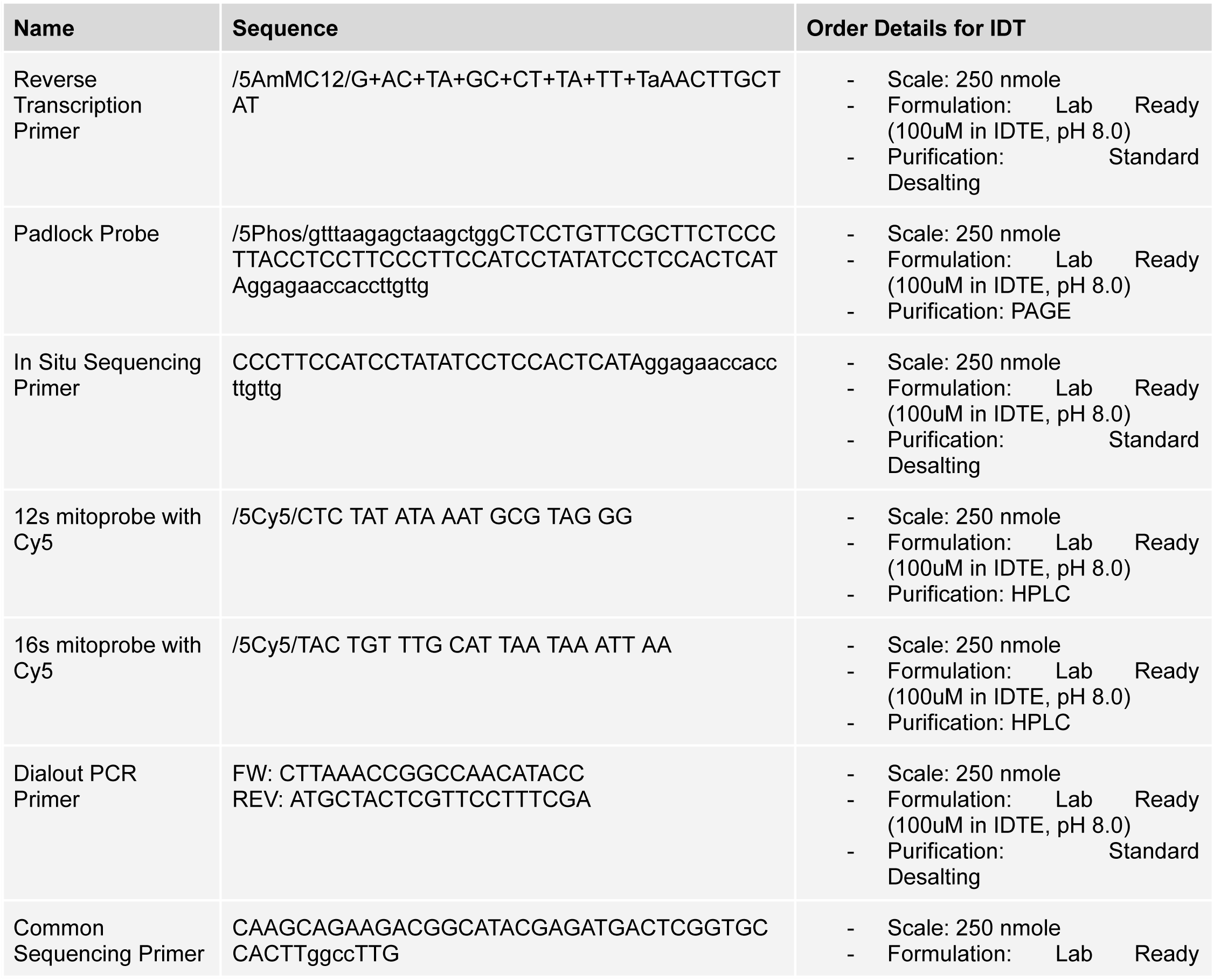
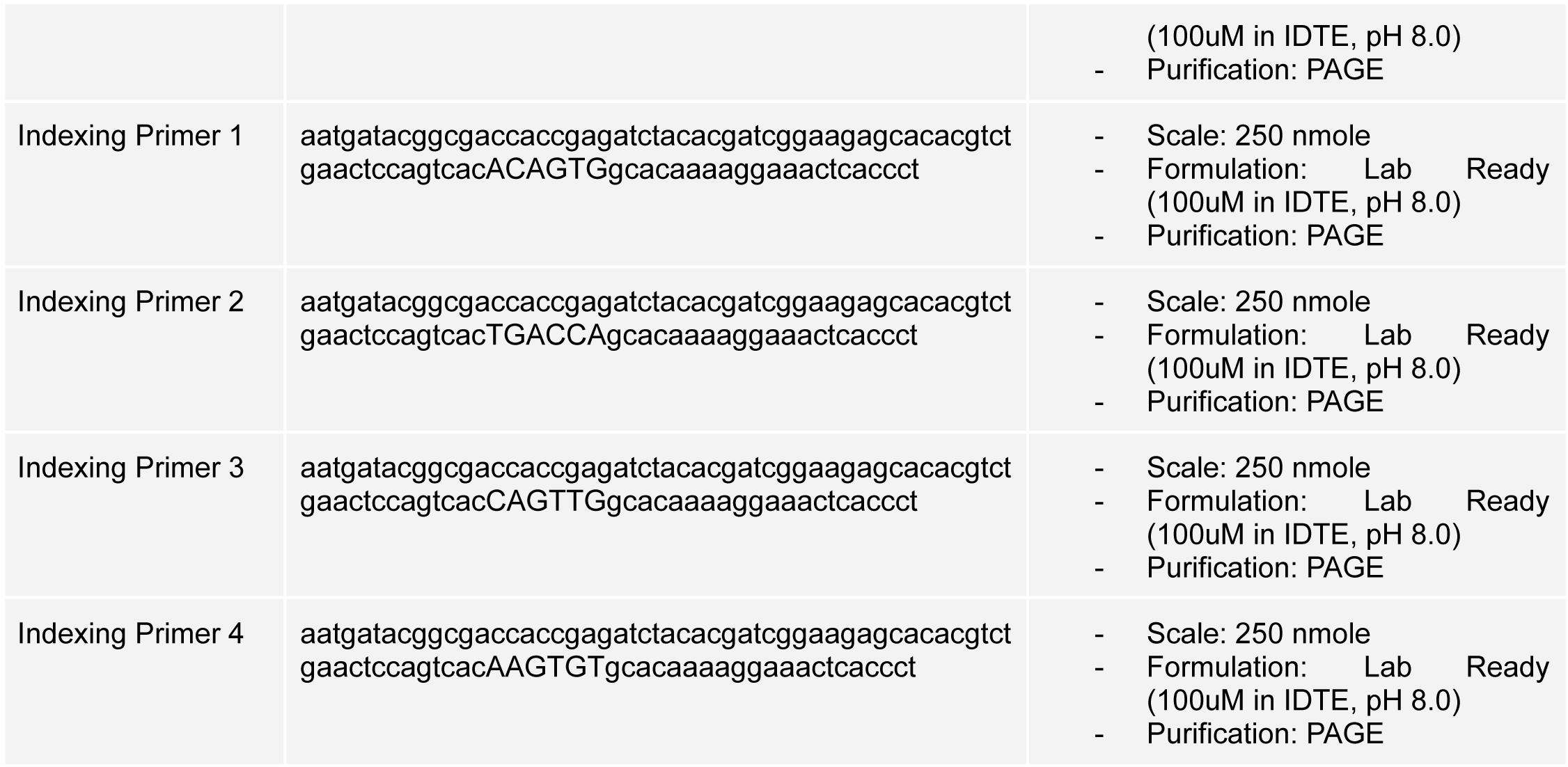
table of oligos used, their sequences and general ordering instructions.

The finalized 5-stain morphological assay panel includes Hoechst, concanavalin A (ConA), wheat germ agglutinin (WGA), phalloidin, and mitoprobe, which stain the nucleus, endoplasmic reticulum (ER), membranes, actin, and mitochondria respectively. This leaves one vacant channel for flexible use, such as a hypothesis-specific biomarker. After imaging cells, ISS was conducted to identify the sgRNA within each cell (Fig. 1d, Methods).

We created a ML enabled data pipeline for high-throughput processing of phenotypic imaging and sequencing by synthesis (SBS) acquisitions in order to create a [morphological feature] * [genetic perturbation] matrix for downstream analysis (Fig. 2a). The process utilized Hoechst staining from all acquisitions to create registration tables, ensuring all morphological phenotyping and SBS acquisitions were spatially aligned and removing the need for manual coarse alignment of field of views during image acquisition, simplifying the lab process. Our approach also includes an improved methodology for base-calling by training a 3-layer fully convolutional neural network (FCN) (Fig. 2a), which increases the deconvoluted cell recovery rate from 66.6% to 78.8% (Fig. 2b), and the determined sgRNA sequences were then matched to the original library design, with ∼2.0, ∼4.0, and ∼4.0 median amplicons detected per amplicon-presenting cell across three separate screens, respectively (Fig. 2c). Samples were sequenced beyond the minimum number of nucleotides needed to call a specific sgRNA within the library. We targeted a minimum Hamming distance of 3 between any two sgRNAs, enabling sgRNA correction in the event of single-nucleotide mismatch. Across the 3 screens, this recovered an additional 5.9%, 5.3%, and 34.8% of cells respectively (Fig. 2d), making the ISS workflow more robust against occasional low-quality cycles or mistakes in base calling. We developed a simple metric for measuring the fidelity of POSH dots using the signal-to-total-ratio (STR, Methods), and found the quality of ISS to be consistently high across cycles of sequencing and across experiments in general (Fig. S2a-d). A single-cell dataset is generated by cropping cell image tiles centered on each nucleus, masked by its corresponding cell mask and associated with a sgRNA identity based on the mapped barcode locations (Fig. 2a). Morphological features are determined using both classical feature extraction and self-supervised learning described below.

**Figure 2.**
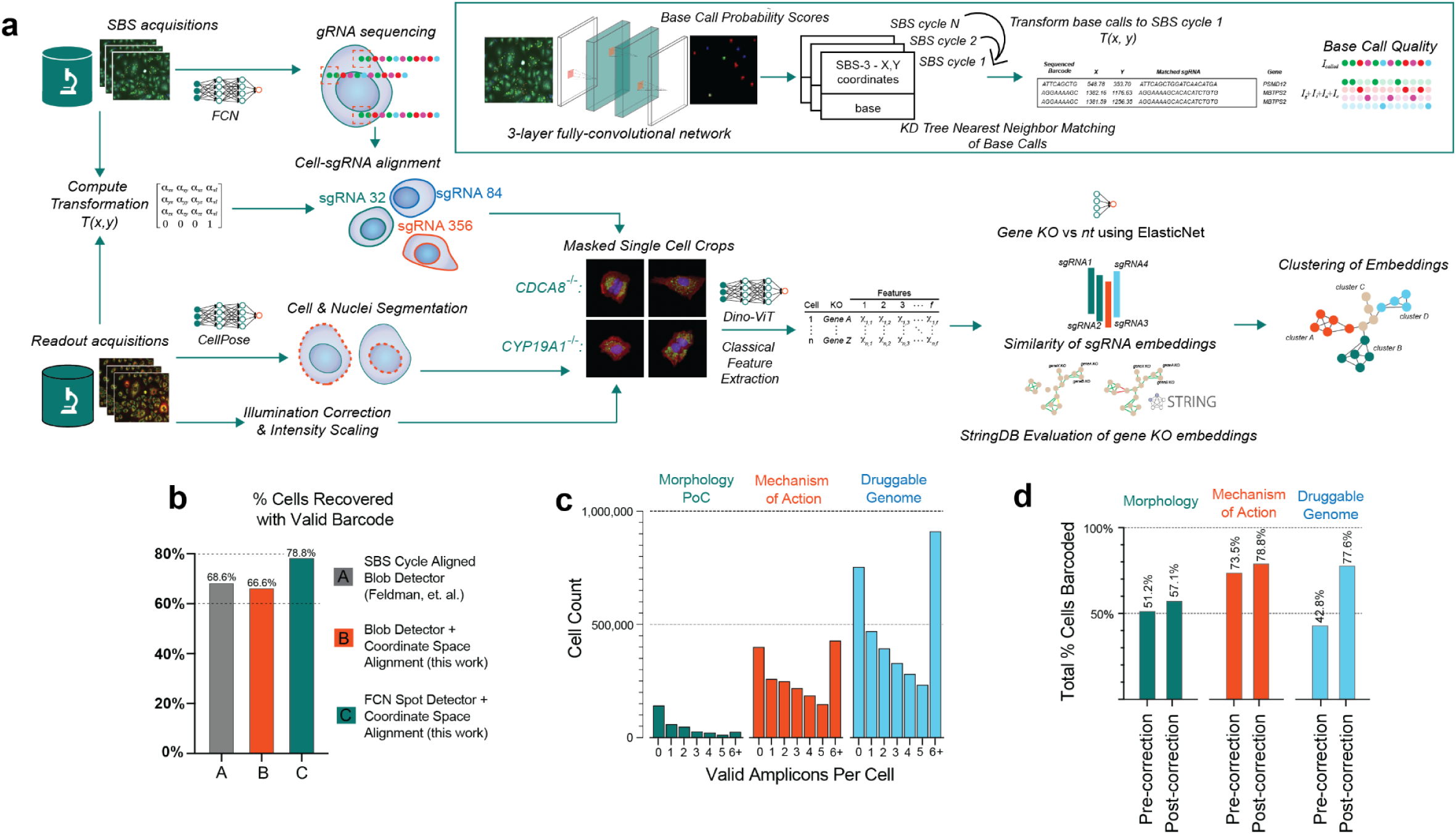
Computational pipeline and QC of POSH platform. **a.** a general overview of the computational pipeline for converting raw phenotyping and SBS images into usable tiles and feature matrices. Sequencing by synthesis (SBS) and readout images are registered using Hoechst staining, followed by amplicon base calling and alignment, in parallel with illumination processing, cell segmentation, and tiling. Multiple analysis methods can then be implemented such as Deep Learning-based methods or direct featurization, followed by embeddings and gene correlation calling. **b.** % cells recovered with a valid sgRNA barcode from Feldman et al^12^ (grey), our study using classical blob detector (orange), and our data using ML (green). **c**. number of valid amplicons per cell across the three screens. Most cells contain at least one valid amplicon. **d.** improvement of cell count based on Hamming correction of miscalled sgRNAs.

### Proof-of-Concept (PoC) with Morphological Regulators

To demonstrate that high-content imaging using Cell Painting captures rich morphological information and can be used to classify gene functions, we set out to perform a small scale CellPaint-POSH experiment against 124 genes with known morphological impact ^32^. A sgRNA library was synthesized to target six key pathways: mitochondrial translation (25 genes), proteasome (27 genes), actin/kinesin (23 genes), unfolded protein response (10 genes), golgi-ER retrograde processes (13 genes), and microtubule/dynein (19 genes), along with 7 miscellaneous genes and 124 nontargeting/intergenic control sgRNAs (Fig. 3a, Supp Data 1), at a representation of 10 sgRNAs per gene for robust analysis. A549 cells were processed with CellPaint-POSH protocol with the aforementioned staining panel. We first assess the fitness effect to validate gene knockouts (KOs). As expected, DepMap-determined common essential genes ^33^ like *COPA*, *KIF11*, and *HSPA5*, as well as several of the proteasome genes, had lower representation than other sgRNAs (Fig. S3a); this provides validation that the KOs are effective and that sgRNAs are correctly sequenced and decoded.

**Figure 3.**
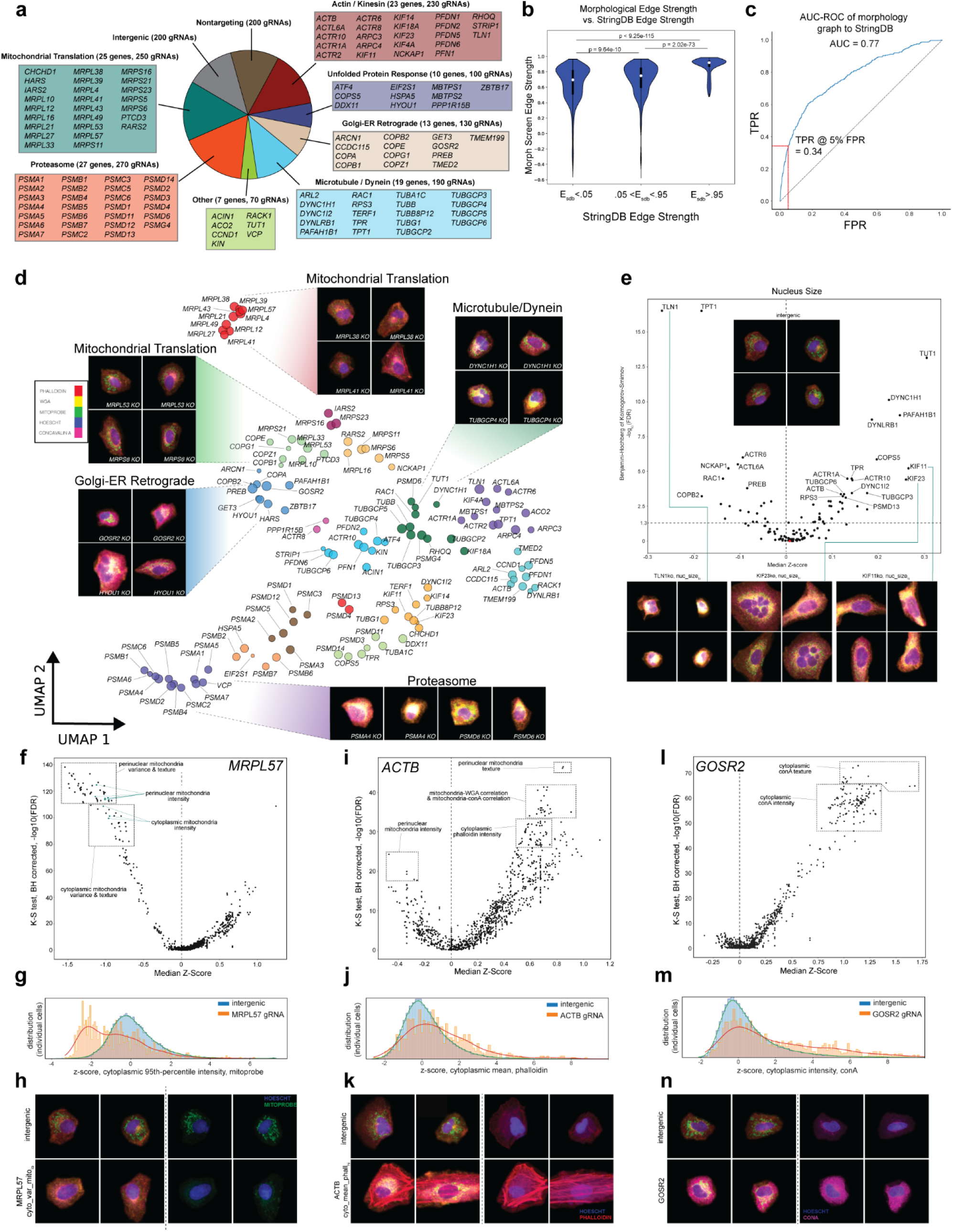
Design and classical image feature based analysis of 124-gene PoC screen. **a**. summary of genes targeted in the screen in order to create distinct morphological effects. **b**. comparison of morphology network gene-gene edges to those from StringDB. Edges defined as strong by StringDB show similarly high edge strengths in our morphological network. **c**. AUC-ROC of CellPaint POSH screen using StringDB network as ground truth, showing strong overlap (AUC=0.77). **d.** UMAP projection of CellStats features showing genes (filtered by AUC > 0.55) clustered by their biological functions. Colors represent Leiden communities, node size represents the median similarity of aggregate sgRNA embeddings targeting the same gene. **e**. volcano plot to identify genes that show significant variation in nucleus size compared to intergenic control sgRNAs. Insets show example tile images of the nuc_size_hi_ and nuc_size_lo_ cells that led to the overall disruption in the distribution of nuclear sizes for their respective knockout populations. Intergenic control in orange. **f**. volcano plot of features comparing *MRPL57* KOs to intergenic controls. **g**. histogram of the *cytoplasmic 95th-percentile intensity* feature in intergenic-sgRNA-presenting and *MRPL57*-sgRNA-presenting cells. **h**. example images of CellPaint on intergenic and *MRPL57* KO cells. **i**. volcano plot of features comparing *ACTB* knockouts to intergenic controls. **j**. distribution of mean cytoplasmic phalloidin intensity of control and *ACTB* KO cells. **k**. example images of the cells with high cytoplasmic phalloidin staining leading to the skewed distribution shown in (j). **l**. volcano plot of features comparing *GOSR2* knockouts to intergenic controls. **m**. distribution of perinuclear conA intensity of intergenic and *GOSR2* KO cells. **n**. example images of the cells with high perinuclear conA intensity leading to the skewed distribution shown in (m).

In total, 163,090 cells were analyzed using a classical image featurization engine similar to CellProfiler ^34^ that extracted 1,301 morphological features in four broad categories: (i) localized pixel intensity statistics, (ii) geometric features of cell segmentation masks, (iii) features characterizing textures that emerge from the different types of staining, and (iv) correlations of intensities across multiple channels (Supp Data 2, Methods). We term this classical featurization engine “CellStats.” Each unique sgRNA was assigned a ’morphology signature’ based on the mean features of all cells containing that sgRNA (Methods). Cosine similarity was determined between these averaged feature vectors to assign similarity scores among all sgRNAs. Similarities between sgRNAs targeting the same gene were specifically compared (Fig. S3b), with a high similarity indicating effective sgRNA cutting and strength of a KO’s morphological phenotype. sgRNA signatures were then further aggregated to the gene level. To systematically assess the accuracy of the biological network emerging from our morphological analysis against prior literature, we compared it to StringDB ^35^, which constructs a gene-gene interaction network in which interactions are weighted by a combination of literature scrubbing, gene-gene interaction databases, co-expression analysis, and organism transfer, and used this as a ground truth comparator. We first compared low, medium, and strong edges in our network to those formed by StringDB and found a very significant increase in correlation between networks at higher StringDB cutoff score (Fig. 3b). We additionally determined the area under the curve of the receiver operating characteristic (AUC ROC) using StringDB edge > 0.95 to define true positives, resulting in a 0.77 AUC, indicating strong overlap with the established network for the same genes (Fig. 3c). We visualize the gene embeddings using Uniform Manifold Approximation and Projection (UMAP) ^36^ grouped by Leiden community detection^37^ to further illustrate the core clusters, notably recreating functional clusters corresponding to the golgi / ER, proteasome, mitochondrial translation (2 distinct clusters), and microtubule / dynein classes (Fig. 3d). Comparison with StringDB and visual inspection of gene clustering both demonstrate that CellPaint-POSH can reconstruct known gene-gene interaction networks.

We took two approaches to further investigate the morphological features driving the gene clustering. First, we identify genes affecting a given morphological feature, such as nuclei size, by computing each KO’s Z-score for this feature (Fig. 3e). Genetic drivers of enlarged nuclei size included gene KOs that yielded strikingly large nuclei (e.g. *KIF11*), or multiple nuclei within the same cell (e.g. *KIF23*), which are consistent with nuclear size, mitotic arrest and spindle polarity phenotypes reported on the same genes in Funk et al^13^. We also observed a significantly smaller nuclear size in *TLN1* KO cells, likely linked to the inability of the cells to form adhesion with their substrate or neighboring cells due to lack of talin-1 protein. Other strong modifiers of nucleus size tended to fall within the Microtubule / Dynein subgroup or encode for members of the actin-related protein (Arp) family, which form key chromatin-remodeling complexes ^38^.

In the second analysis approach, which we term “differential morphology analysis,” we examine all CellStats features modulated by a particular genetic perturbation, relative to the control cells. Differential morphological features are similar in concept to differentially expressed gene analysis commonly used in comparing two sets of transcriptomics data. As an example, we selected one gene that represented a single point in one of the tightly-connected clusters within the network (*MRPL57*, Fig. 3d). We conducted Kolmogorov-Smirnov (K-S) tests between *MRPL57* cell features and intergenic-targeting sgRNA controls, and found that perinuclear mitochondrial intensity, variance, and texture were the most significantly different features of *MRPL57* KO (Fig. 3f-g), which matches the mitochondrial dysfunction anticipated with the KO. Representative plots of *MRPL57* KO cells driving these mitochondrial scores depict the low-mitochondria phenotype (Fig. 3h). We next examined cytoskeletal protein *ACTB*. Interestingly, phalloidin scoring indicated a significant increase in actin presence within the cells; this provides evidence for compensatory up-regulation of other actin isoforms upon loss of β-actin ^39^ (Fig. 3i-k). Lastly we examine Golgi SNAP Receptor Complex Member 2 (*GOSR2)*, whose loss of function likely leads to progressive myoclonus epilepsy in patients ^40^. Examining the differential morphological features upon *GOSR2* KO (Fig. 3l), its unique morphological phenotype was driven largely by increased intensity, variance, and texture of Golgi/ER specific ConA staining (Fig. 3m-n), matching its likely biological mechanism ^41^, and providing a disease phenotype that could be adapted into a disease-modifier screen.

### Mapping MoA with Self-Supervised Vision Transformer Learning

Encouraged by our morphology-driven PoC, and inspired by previous efforts that use Cell Painting information to cluster compounds by their MoA ^24,27^, we next attempted to assess the performance of CellPaint-POSH at phenotypic clustering of genes by their annotated mechanism of action, irrespective of previous reported morphological phenotypes. We curated a list of 300 genes whose gene products are targeted by tool compounds with well annotated MoAs ^42–44^ (Supp Data 3). We built self-supervised vision transformer (DINO-ViT) models ^45^ that can extract meaningful image representation with no labels, and compared between the following imaging representation techniques: 1) classical morphological featurization (“CellStats”); 2) self-supervised DINO-ViT model, trained on ImageNet ^46^ containing ∼1.2 million natural image data: e.g. animals, vehicles, and tools (“ImageNet-dino”); 3) DINO-ViT embeddings trained on ∼1.5M single cell Cell Painting images from the 300-gene MoA experiment (“CP-DINO 300”, Methods, full cell tile image dataset provided in Supp Data 4). All three featurization methods are capable of morphologically classifying many genetic perturbations from non-targeting control cells (Fig. 4a-b), including ImageNet-dino which is not trained on cellular morphology data. This is consistent with previous reports of transfer learning ^47^. Nevertheless, CP-DINO 300 trained on bioimaging data yielded a more informative embedding that has higher median prediction accuracy than the other two models (Fig. S4a-b), and correctly classified more perturbations with better accuracy (Fig. 4c). CP-DINO 300 also recovered more known biological relationships from StringDB as measured by cosine similarity of the aggregate gene KO embeddings (Methods) than the other two models (Fig. 4d).

**Figure 4.**
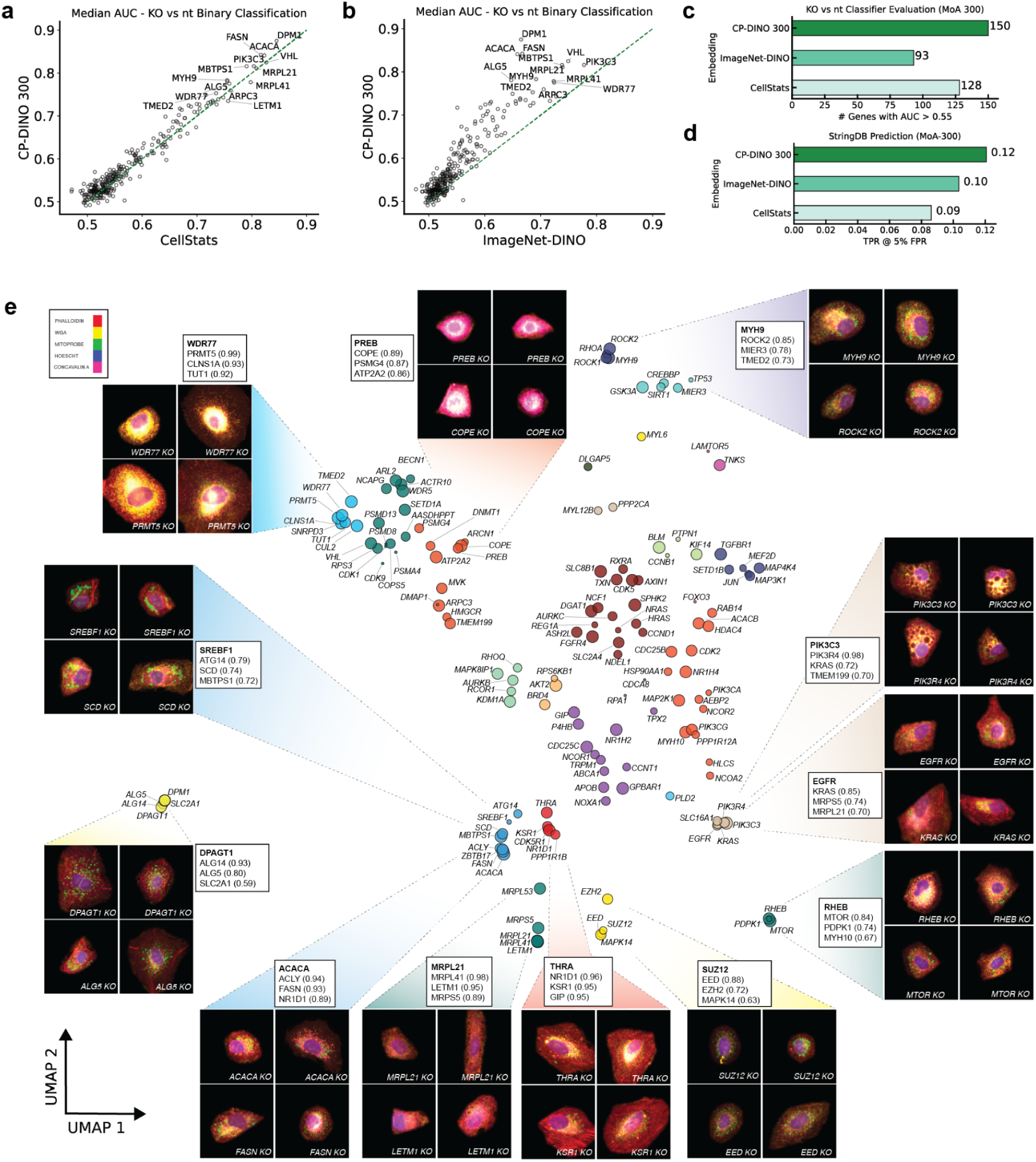
DINO trained embedding outperforms Cellstats and ImageNet-trained embedding. **a-b.** Comparison of feature embedding methodologies based on median AUC of binary classification of KO from WT for each genetic perturbation. **a.** CP-DINO 300 vs. CellStats **b.** CP-DINO 300 vs. ImageNet-dino. **c.** Number of genetic perturbations classified at AUC > 0.55 by different representation models. **d.** True positive StringDB edges predicted (at 5% false positive rate) performance using gene-gene similarity calculated from different representation models. **e.** UMAP projection of DINO-ViT features showing genes (filtered by AUC > 0.55) clustered by their cellular component localization. Colors represent Leiden communities, node size represents the median similarity of aggregate sgRNA embeddings targeting the same gene. Cell tile images and top neighbors (cosine similarity in parenthesis) of select KOs are shown.

Similar to CellStats, CP-DINO 300 representation of the Cell Painting assay allows reconstruction of gene networks, and the identification of new pathway components in a hypothesis-free way using Leiden community detection (Fig. 4e) ^37^. Specifically, we were able to reconstruct the genetic modifiers of glycoprotein biosynthesis, mitochondrial translation, actin/cytoskeletal organization, lipid metabolism, *PI3K*/*Akt* activation, mTORC1 signaling, PRC2 complex, where morphological similarity associates genes from the same pathways / protein complexes. For example, genes with the highest similarities to *SUZ12* included the other two PRC2 complex members *EZH2* and *EED* ^48^ (Fig. 4e); genes similar to mTORC1 activator *RHEB* included other pathways activators *mTOR* and *PDPK1* ^49^. Interestingly, the network was capable of clustering key components of the lipogenesis pathways without the use of an explicit lipid stain, including the core fatty acid synthesis enzymes (*ACLY:* ATP Citrate Lyase*, ACACA*: Acetyl-CoA Carboxylase Alpha*, SCD*: Stearoyl-CoA Desaturase, and *FASN*: Fatty Acid Synthase), and upstream AKT signaling regulators (*PIK3C3:* PI3K Regulatory Subunit 3*, PIK3R4*: PI3K Regulatory Subunit 4) that all contribute to lipogenesis ^50^ (Fig. 4e). Representative images of gene KOs with the most distinguished phenotypes (highest binary predictive scores) from each cluster are shown (Fig. 4e).

### Deep Learning Enables Novel Discovery via Druggable Genome Pooled Optical Screen

We next sought to scale CellPaint POSH to the druggable genome to demonstrate the throughput needed for discovery screens, and to further assess the generalizability of Cell Painting across more diverse sets of genes and pathways. To this end, a library of 1640 genes was assembled based on the Tier 1 list of the “Druggable Genome” library ^17^, spiked in with a subset of morphological genes and known mTORC1 pathway genes for controls (Supp Data 5). In addition to Cell Painting, anti-phospho-S6 (pS6) antibody with AlexaFluor 750-conjugated secondary antibody was used in the 6th channel as an established biomarker ^51^ used extensively in mTORC1 studies including genome-wide screens ^22,52,53^. In order to improve throughput and reduce labor-intensive manual work, we built and deployed fully automated liquid handling and microscopy for the ISS workflow (Methods, video demonstration in Supp Data 6). We formally assessed sgRNA library abundances as determined by ISS with that measured by genomic DNA next-gen sequencing (NGS), by comparing the enrichment and depletion of sgRNAs targeting fitness genes. We again re-discovered known essential genes ^33^, and the measured fitness effects were comparable between POSH and NGS (Fig. S5a-c), indicating that there is little detection bias in POSH ISS.

We explored if the added diversity in the 1640-Gene druggable genome POSH dataset would lead to better self-supervised image representation. To that end we trained CP-DINO 1640, and found that while it performs similarly to CP-DINO 300 at binary classification of genetic KOs vs. negative controls in the 1640 dataset (Fig. 5a-b), CP-DINO 1640 captures more semantically meaningful structure in the data as demonstrated by its more accurate predictions of StringDB gene-gene interactions than CP-DINO 300, ImageNet-dino, or CellStats (Fig. 5c).

**Figure 5.**
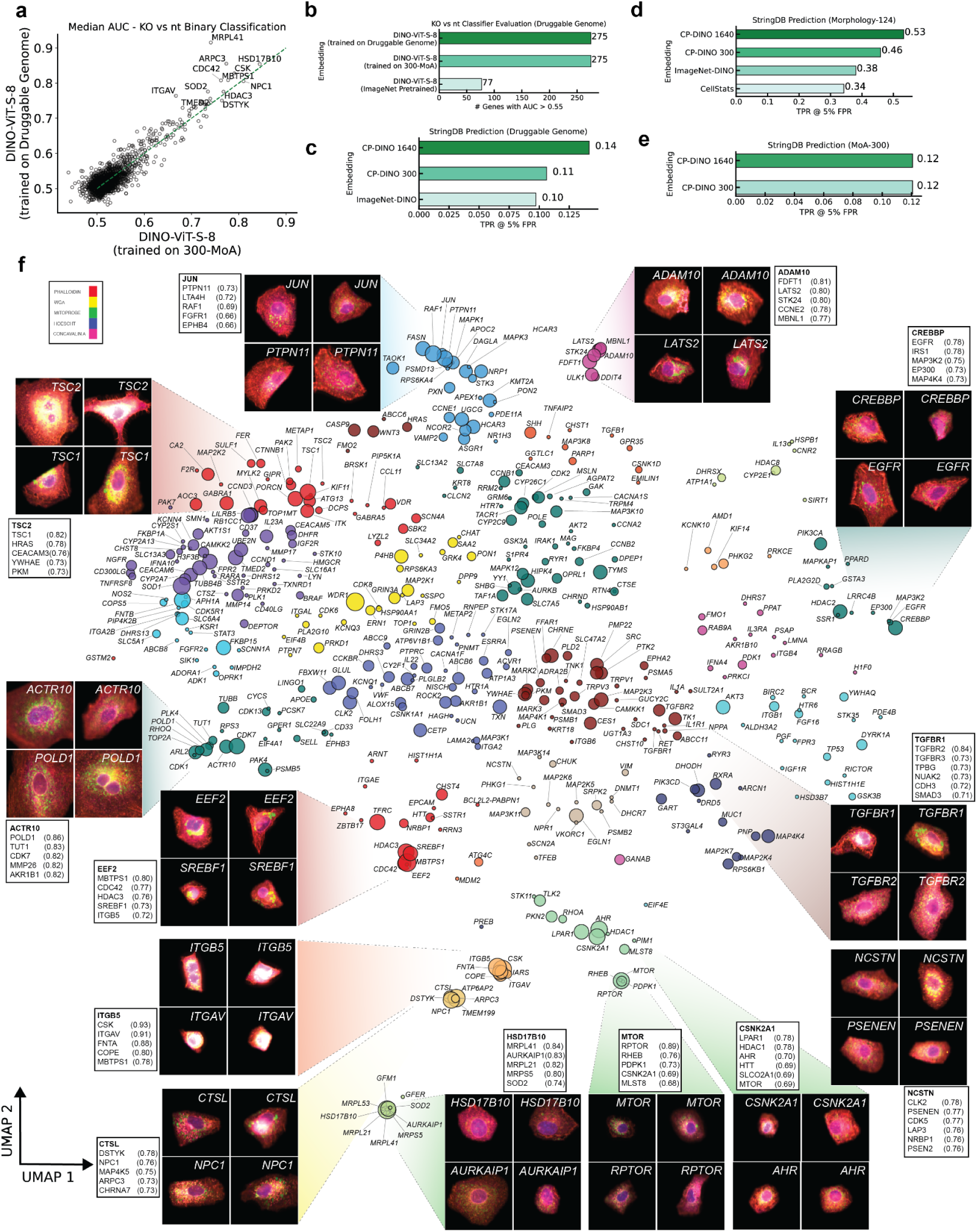
Morphological analysis of druggable genome discovery screen. **a**. Comparison of feature embedding methodologies (CP-DINO 1640 vs. CP-DINO 300) based on median AUC of binary classification of KO from WT for each genetic perturbation. **b.** Number of genetic perturbations classified at AUC > 0.55 by different representation models. **c.** True positive StringDB edges predicted (at 5% false positive rate) from 1640 druggable genome dataset using gene-gene similarity calculated from different representation models. **d.** True positive StringDB edges predicted (at 5% false positive rate) from a held-out 124 gene PoC dataset using gene-gene similarity calculated from different representation models. **e.** True positive StringDB edges predicted (at 5% false positive rate) from 300 gene MoA dataset using gene-gene similarity calculated from different representation models. **f.** UMAP projection of DINO-ViT features showing genes (filtered by AUC > 0.54) clustered by their biological functions. Colors represent Leiden communities, node size represents the median similarity of aggregate sgRNA embeddings targeting the same gene. Cell tile images and top neighbors (cosine similarity in parenthesis) of select KOs are shown.

We next evaluated all image representation models on held out data: the 124-gene morphology PoC perturbation dataset. While all 3 self-supervised DINO-ViT models generalized out of experiment and outperform CellStats at predicting StringDB networks (Fig. 5d), CP-DINO 1640 in particular surpassed prediction accuracy from CP-DINO 300 and ImageNet-dino (Fig. 5d), and faithfully reconstructs gene networks (Fig. S5) by Leiden clustering. The superior performance of CP-DINO 1640 is unlikely a result of trivial memorization, as the 1640-genes druggable genome library and 300-genes MoA library share similar numbers of overlapping genes with the 124 PoC library (30 and 26 genes respectively). Lastly, when evaluated on the 300-MoA dataset, both CP-1640 and CP-300 performed similarly in StringDB prediction (Fig. 5e), likely because the latter already learned the full variance in the corresponding dataset.

We constructed gene networks from the druggable genome dataset using CP-DINO 1640 model and Leiden community detection (Fig. 5f). Overall, many genetic pathways and known protein complexes emerged from network analysis of imaging features from the unbiased morphological profiling. As an example, a cluster of morphologically-related genes formed by *MAP4K4*, *MAP3K2*, *PIK3CA*, *EGFR* and *CREBBP* / *EP300*, all known regulators of EGF signaling signaling ^54–57^. As another example, top related genes to *TGFBR1* included other known TGFb genes such as *TGFBR2*, *TGFBR3* and *SMAD3* (Fig. 6d)^58^. Among the top genes similar to *NCSTN* is *PSENEN*, both of which encode for the gamma-secretase complex components nicastrin and presenilin enhancer 2, respectively ^59^. A cluster with strongest gene-gene similarities include *ACTR10*, *POLD1*, *TUT1*, *CDK7*, *TUBB*, and *TOP2A*. Interestingly, while their KO phenotypes are similar, these genes perform different biological functions (e.g. *TUT1* encodes a nucleotidyl transferase that functions as both a terminal uridylyltransferase and a nuclear poly(A) polymerase, while *PLK4* is involved in centriole duplication). We note that most of these are reported essential genes ^33^ and confirmed by sgRNA abundances in our experiment (Fig. S5d). Indeed, *PLK4* is a known essential regulator of cell cycle ^60^, and it is possible that nucleotidyl transferase *TUT1* may also play an important role in cell division. Consistent with this hypothesis, CellStats analysis suggested that *TUT1* KO induces large nuclei size (Fig. 3e). In addition, Luke et al. recently reported both *TUT1* and *PLK4* KOs produce larger cell area, nucleus area and nuclear DNA integrated intensity, consistent with the phenotype of cell cycle regulator genes such as CDC proteins ^13^. Both Deep Learning (DL) and CellStats analysis nominate the potential role of *TUT1* in cell cycle regulation.

**Figure 6.**
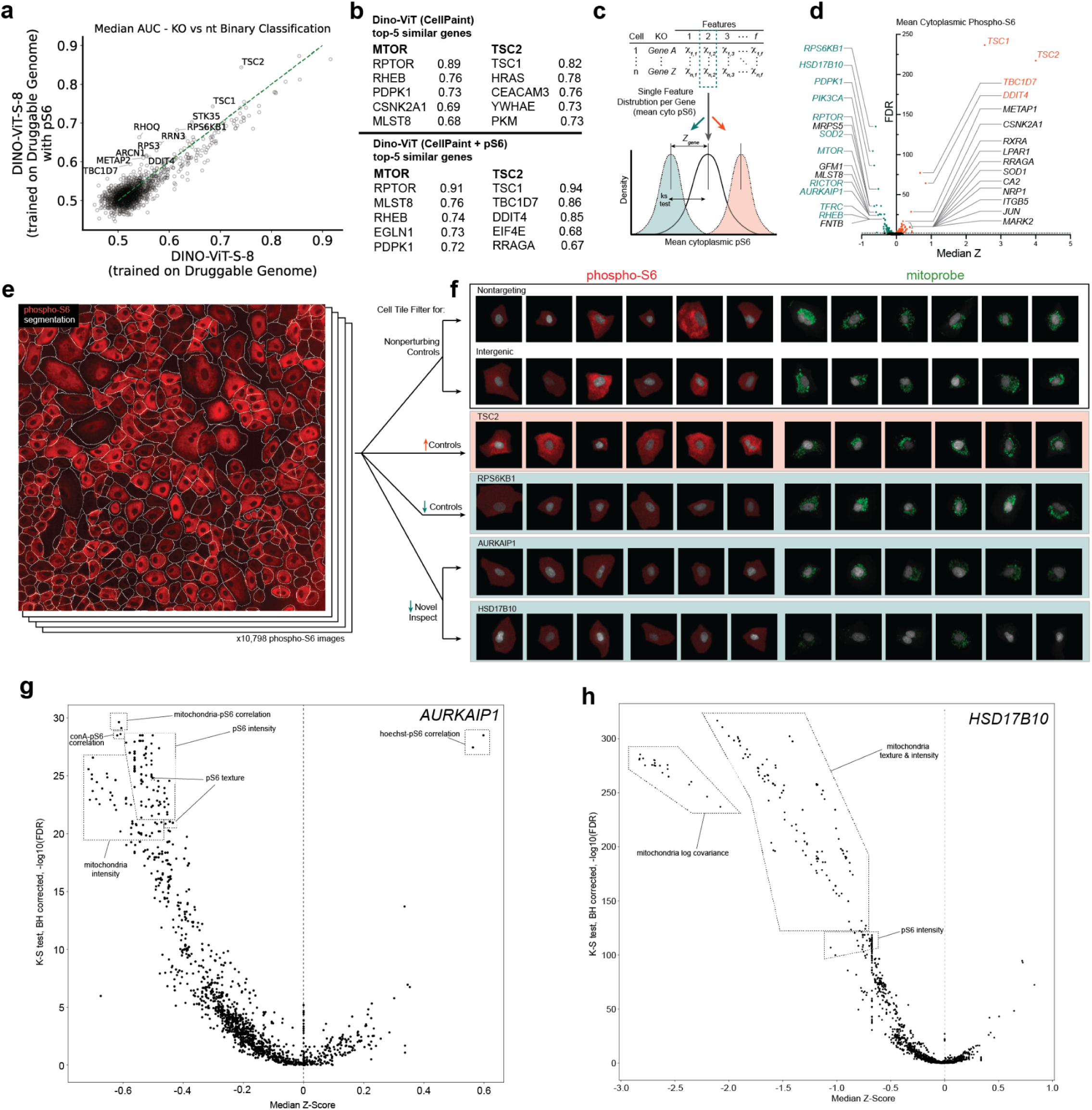
Combining morphology and pS6 biomarker analysis for mTORC1 pathway discovery and validation. **a**. Comparison of feature embedding methodologies (CP-DINO 1640 with and without pS6 information) based on median AUC of binary classification of KO from WT for each genetic perturbation. **b**. Top neighbor genes to mTOR and TSC2 (mTORC1 activator and inhibitor respectively) predicted with and without pS6 are shown. Numbers indicate cosine similarity. **c**. the pipeline for conducting single-cell based analysis via Kolmogorov-Smirnov 2-sided, 2-tailed statistical test with benjamini-hochberg correction, in contrast to methods used for flow-based screens such as MAUDE (Fig. S7a). **d**. results of single-cell based, one-feature analysis of cytoplasmic phospho-S6. Teal genes are well established mTOR upregulators, orange genes are well established mTOR downregulators, black genes are potentially novel pS6 modifiers. **e**. representative pS6 field with segmentation. **f**. example pS6 and mitoprobe staining images from control, known, and potentially novel regulators of mTORC1. **g-h**. differential feature analysis on *AURKAIP1* and *HSD17B10*.

Another major pathway that emerged from gene network analysis is mTORC1, where 3 known mTORC1 complex proteins form a tight cluster: *RPTOR*, *mLST8*, *mTOR* (Fig. 5f) ^61^. Other members of the core mTORC1 cluster include well established activators of the PI3K/AKT/mTOR pathway such as *CSNK2A1* and *PDPK1* ^62,63^. In contrast, mTORC2 core complex component, *RICTOR*, clusters away from this group, likely because of its primary role in a distinct *mTOR* complex with different functions and morphological consequences on the cell. In addition, mTORC1 inhibitors *TSC1* and *TSC2*, whose loss of function (LoF) mutations lead to the eponymous genetic disorder ^64^, cluster away from mTORC1 activators as expected (Fig. 5e).

Given the clustering of mTORC1 regulators, we asked if the inclusion of the pathway biomarker pS6 may improve the discovery of relevant genetic perturbations. We thus trained CP-DINO models with Cell Painting + pS6 channel information, and found that while both CP-DINO 1640 and the new CP-pS6-dino 1640 have similar sensitivity in predicting genetic perturbations, pS6 notably improves the detection of many mTORC1 inhibitors (*TSC1*, *TSC2*, *DDIT4*, *TBC1D7*) (Fig. 6a). Indeed, while mTORC1 activators (defined as top genes morphologically similar to mTOR) are similarly predicted with or without pS6 (Fig. 6b), interestingly mTORC1 downregulators (defined as genes morphologically similar to *TSC2*) include relevant hits such as *TBC1D7* and *DDIT4* only when pS6 information is taken into model building ^65,66^. In order to further validate morphologically discovered mTORC1 regulators against the ground truth biomarker, we systematically analyzed genes that influence pS6 intensity with two methods: 1) a binning approach, grouping cells with similar levels of pS6 intensity changes, simulating enriched bins of cells produced by FACS-based CRISPR screens (Fig. S7a); 2) a distribution based approach, taking full advantage of the single cell resolution of our data (Fig. 6c, Methods). Both analyses resulted in comparable hits, effectively identifying many of the established regulators of the mTORC1 pathway ^22,52^ (Fig. 6d, Fig. S7b). Most top mTORC1 activators as defined by morphological similarity to mTOR, including *mTOR*, *PDPK1*, *RPTOR*, *mLST8*, *CSKN2A1* mentioned above (Fig. 6b), are also top regulators of the pS6 biomarker (Fig. 6d). Interestingly, mTORC1 hits discovered by the pS6 assay can be further stratified into different subclusters by generic morphological analysis (Fig. 5e). For example, certain pS6 hits *GFM1*, *HSD17B10*, *MRPS5*, *MRPL21*, and *AURKAIP1* form a morphological cluster that is separate from other pS6 hits and core mTORC1 regulators (e.g. *mTOR*, *PDPK1*, and *RPTOR*) (Fig. 5e), suggesting that the former group of genes may regulate mTOR biology through a distinct mechanism. Based on their biological functions ^67–69^, we hypothesize that mitochondrial translation and metabolism may be involved in mTORC1 regulation. While the role of mTORC1 in regulating mitochondrial function has been reported^70^, the reverse causation is not fully understood. We performed differential feature analysis on these genes, and showed that genes such as *AURKAIP1* and *HSD17B10* not only lower pS6 levels in the cell but also have a significant impact on mitochondrial staining (Fig. 6e-f). Furthermore, CellStats based differential feature analysis shows that pS6 and mitochondria-related morphological features are among the top changes upon *AURKAIP1*, *HSD17B10*, and *MRPS5* KOs (Fig. 6g-h, Fig. S7c). Consistent with our findings, *HSD17B10* was recently discovered as a novel mTORC1 regulator in a genome-wide screen against pS6 ^22^. In addition, mitochondrial translation defects caused by the deletion of another mitochondrial elongation factor *mtEF4* in *C. elegans* and mouse have been shown to regulate *mTOR* and cytoplasmic translation ^71,72^ as a compensatory mechanism to cope with mitochondrial stress. We hypothesize that other mitochondrial genes like *AURKAIP1* and *HSD17B10* may have an effect on mTORC1 in our experiment. Interestingly, mutations in *HSD17B10* have been linked to recurrent seizures and intellectual disability ^73,74^, two symptoms also commonly observed in mTORopathies ^75^.

## Discussion

### Pooled morphology CRISPR screening for hypothesis-free exploration

Pooled CRISPR screens have emerged as a powerful technique to interrogate multiple genetic interventions in parallel; they are considerably more cost effective and scalable than arrayed screens, and are able to greatly ameliorate batch effects. CellPaint-POSH combines the benefits of pooled CRISPR screening with the rich cell state information provided by Cell Painting. While biomarker based screens can enumerate pathway regulators along a unidimensional axis (e.g. pS6 intensity), morphology analysis can identify rich structures spanning multiple biologies. Indeed, across 3 experiments, morphological data clustered genes that affect the same cellular components (e.g. proteasome, golgi/ER, mitochondria inner membrane), molecular functions (e.g. protein glycosylation, cytoskeletal organization, fatty acid synthetic process, chromatin modification) or biological pathways (e.g. EGF, mTOR, TGFb), and was able to do so without prior hypothesis or explicit biomarkers/assays. Morphological guilt-by-association analysis also helps generate novel hypotheses, for example linking uridylyltransferase to cell cycle regulation. In the same example, we note that functionally distinct genes (e.g. *PLK4* and *TUT1*) may have similar morphological phenotypes when deleted, suggesting shared functions for genes that are not known to be related.

We also expand the CellPaint panel with an additional channel, and show that the resulting “CellPaint + 1” design enables the use of a biomarker (e.g. pS6) that provides built-in validation of morphological discoveries from the same experiment. We demonstrate that the rich morphology readout can stratify genes that manifest similarly in biomarker studies (e.g. pS6) into sub-clusters representing molecular mechanisms (e.g. core mTORC1 v.s. mitochondria translation), demonstrating the benefit of coupling high dimensional data with ML analysis. We envision that our hypothesis-free approach can be a useful tool for disease modeling and drug target discovery, particularly when the disease biological pathway or biomarker is not completely understood, similar to recent applications of perturb-seq in disease research ^76^. Techniques that further increase the dimensionality of cellular imaging, such as multiplexed protein / RNA detection ^77,78^, may provide finer grained and more diverse phenotypic readouts and reveal additional biological insights.

Pooled optical screening does have some important technical limitations. First, pooled optical screening, like any other high-content image-based assay, can be sensitive to imaging artifacts such as poor focus and imaging channel bleedthrough, requiring careful image QC and staining panel design. Second, ISS signals can be low in certain cell-types with lower expression levels of sgRNA spacer containing transcripts, requiring higher cell input to compensate for lower data recovery (data not shown). Third, pooled optical screening requires instance (per cell) segmentation, which is difficult in cell-types that have overlapping structures (e.g. neurons with extensive and long dendrites, hepatocytes that grow in 3D, data not shown). We have shown that our use of deep-learning assisted cell segmentation helps ameliorate some of these challenges, but further development along those lines will improve the analysis for these more challenging cell types. Fourth, while we demonstrate that high-dimensional screening provides strong phenotypes that cluster effectively, the interpretation of these phenotypes can be more challenging than in transcriptomics. Leveraging both DL analysis for better image representation and classical image analysis for interpretability may be a good combination strategy, as a way to explain why *TUT1* may be related to cell cycle regulators for example.

During the final preparation of this manuscript, a similar method (PERISCOPE) emerged that combined Cell Painting and pooled optical screening to build an unbiased morphological atlas ^79^. Our work is differentiated in a number of ways that enable the broad applicability and scaling of our approach. On the experimental side, we have addressed the spectral overlap challenges between the CellPaint panel and the ISS barcodes by using a novel, structural biology inspired RNA-FISH-probe for mitochondrial staining, avoiding the need for custom-conjugation probes. We also developed a closed-loop automation system for in situ sequencing, which was successfully utilized in our druggable-genome discovery screen (Supp Data 6); this capability is critical for scaling throughput and ensuring process and data robustness. Computationally, our use of ML enabled significant improvements to many critical steps during POSH analysis, including base calling, cell segmentation and image representation. For base calling, our approach using a 3-layer convolutional network followed by global registration and transformation of coordinates can be executed without manual alignment of field-of-view images at acquisition time and without manual parameter tuning, yielding improved sensitivity and a simplified lab workflow. Moreover, the use of ML for cell segmentation and guide calling considerably enhanced our ability to utilize more of the cells and hence greatly increased the throughput of our platform. Finally, we note that while we have focused our presentation on our work in A549 cells, we have also successfully applied CellPaint-POSH in HepG2s, as well as in iPSC-derived cortical neurons, motor neurons and stellate cells (data not shown); we believe it can be adapted to most cell types that could be grown and transduced in vitro.

### ML-based analytical methods

Morphological analysis is heavily influenced by the features extracted, and we rigorously compared classical image feature extraction with self-supervised DL methods. Classical image analysis uses expert engineered features such as intensity, texture, shape and size of cells and their internal compartments. While such an approach is simple and interpretable, it incorporates human bias and does not capture the full extent of biological variability in the dataset. Deep learning is an alternative approach that is less susceptible to human bias, but existing models are trained on natural image datasets and may not align well with the statistical patterns in bioimaging data. Recently, weakly supervised DL and self-supervised auto-encoder methods have been used to learn embedding directly from cellular images ^80,81^. Our work leverages DINO and Vision Transformer (ViT) ^45^, which has been shown to achieve state-of-the-art performance in learning representations from natural images. We demonstrated that self-supervised models with ViT architecture trained on single cell feature embeddings provide better predictive accuracy and improved gene interaction prediction when compared with classical single cell featurization methods and ImageNet pretrained self-supervised models.

The information richness of a high throughput and high dimensionality CellPaint-POSH dataset, combined with the power of deep learning featurization, allows a diverse range of analyses to be conducted based on a single experiment, greatly accelerating biological discovery. For global gene network analysis, we borrowed heavily from standard single-cell RNAseq methods and found that “guilt by association” was one of the most effective ways of identifying known and novel components of pathways. Our work demonstrates that this approach helps identify both known and novel biology via a simple and uniform analysis. As a future direction, other dimensionality reduction techniques that define and explore multi-dimensional latent spaces, similar to singular value decomposition or non-negative matrix factorization used in single-cell RNAseq analysis ^82,83^, may further improve the computational resolution of mapping gene-function relationships and help reveal pleiotropic gene function (such as in the case of *TUT1*).

Importantly, we also showed that the single cell feature representation models trained from one dataset generalized well to an unseen dataset, allowing a pre-trained model to be easily deployable in interpreting future experiments without expensive and time-consuming de-vno training every time. We note that when comparing CP-DINO 1640 with CP-DINO 300, the former captures more semantic structure in the data, even though it is trained with a similarly sized dataset as the latter (both ∼1-1.5million cell tile images). We reason that this is due to the greater degree of phenotypic diversity present in the 1640 dataset thanks to the five-fold more genetic perturbations. We predict that increasing the in-house training data size and (even more importantly) phenotypic variance -e.g. scaling to genome-wide POSH experiments or even experiments done in different cell types - may further improve the DL model and make it more generalizable. In addition, publicly available non-POSH datasets such as the JUMP Cell Painting collection ^84^ can be incorporated into the training data. Notably, the self-supervised ViT architecture requires no labels, and thus can be trained on bioimaging datasets without upfront annotations, circumventing a significant bottleneck and enabling model training on a broad range of internal and external data sets. Thus, over time, we can build richer foundation models that provide an unbiased distillation of the phenotypic landscape of cell morphology. These models can be used to interrogate new gene-phenotype relationships and deliver new insights regarding diverse pathways and biologies, paving the way towards novel biological discoveries.

## Methods

### sgRNA Design, Synthesis and Ordering

Oligo libraries were designed that targeted 1) morphological phenotypes (124 genes, 10 sgRNAs per gene), 2) the druggable genome (1640 genes, 4 genes per sgRNAs) and 3) mechanism of action study (300 genes, 4 sgRNAs per gene). The gene list was then put through a sgRNA selection pipeline that pulls sgRNAs (∼20bps long) from Brunello ^85^ and Toronto KnockOut version 3.0 ^86^ sgRNA repositories. sgRNAs targeting the same region were removed by filtering on a Levenshtein distance of less than 3 and a max subsequence match length of greater than 15. sgRNAs without a BsmBI restriction enzyme cutting site used for golden gate cloning were discarded. The sgRNAs were then sorted by their on-target score per gene as reported if available ^85^. A greedy approach was used to iteratively pick *N* sgRNAs per gene in the ascending order of the number of sgRNA designs available per genes. In order to allow for correction of at least 1 base-pair error throughout the library, we also set a hamming distance cutoff requirement of 3 for selecting any subsequent sgRNA and select the sgRNAs iteratively until we have sgRNAs for all genes in our input design list. Non-targeting and intergenic sgRNAs were then placed in each library as controls. The last step of the pipeline appends the oligo sequence flanking the sgRNA that allows cloning into our backbone, while including the dialout primers needed for library specific amplification. Libraries were duplicated or triplicated before submitting to a vendor for synthesis, based on the size of the initial sgRNA list. This helped to prevent jack potting and propagation of errors while also keeping the relative ratio of controls to targeting sgRNAs the same. We then ordered the library from Twist Bioscience.

### Library Cloning

Library pools received from Twist Bioscience were spun down at 11,000 rcf for 2 mins then resuspended to 5-10ng/mL in IDTE pH8.0 1xTE solution (11-05-01-13, IDT) . We incubated the freshly resuspended library pool for 5 minutes at room temperature then vortexed for 20-30 seconds, then spun down for 2 minutes at 11,000 rcf.

All of our oligo libraries were cloned into a customized lentiviral CROPseq-like vector (plasmid backbone) of which some features include puromycin resistance, TagBFP, padlock sites and Esp3I/BsmBI sites for cloning (Supp Fig 1b). Per library to be cloned, the backbone was digested using 2 ug of plasmid backbone, 2 uL of FastDigest Esp3I (BsmBI) enzyme (ThermoFisher, FD0454), 0.5 uL 100 mMM DTT (Bioworld, 3483-12-3), 2 uL of 10X FastDigest buffer, and molecular biology (molbio) grade water (46-000-CI, Corning) added to a final volume of 50 uL. Solution was then incubated in a thermocycler at 37C for 2 hrs. Then 1 uL of fast alkaline phosphatase (ThermoFisher, EF0652) was added and the sample was incubated at 37C for 45 mins then heat inactivated at 75C for 5 mins. We then followed the E-Gel CloneWell II Agarose Gel protocol to gel-purify the digested backbone. Sample was eluted and DNA was quantified using a Nanodrop. Typical recovery of DNA was 50-70% of the starting solution. Digested vector was then stored at -20C.

A 200uL dialout PCR solution was made in molbio grade water (46-000-CI, Corning) to a final concentration of 1x Q5 PCR (M0492S, NEB), oligo pool (2.5ng final), 1x evagreen (Biotium, 31000T), 0.25 uM dialout primers (see Table 1). qPCR was run on the sample according to the protocol with 1x 98C for 10 seconds > 16x 56C for 30 seconds, 72C for 30 seconds, and 72C for 30 seconds > 1x 72C for 120 seconds. qPCR was run until the last sample entered the exponential phase, up to a maximum of 16 cycles. PCR product was cleaned up using SPRIbeads (Beckman Coulter, B23318) at 0.65x ratio (ex: 32.5uL SPRI beads and 50uL of sample). DNA was eluted typically in 50 uL elution buffer after two 80% ethanol washes. We then quantified the DNA sample using a Qubit 4 Fluorometer (invitrogen, Q33238) and the high sensitivity 1x dsDNA kit (invitrogen, Q33231).

Golden gate assembly was done using the NEB Golden Gate Assembly Kit (BsmBI-v2) E1602 protocol. In short, the NEB reaction was prepared in a PCR strip tube to a final amount of 200 ng of pre-digested CROP-seq backbone, 80 ng of the insert (sgRNA library), 2.5 uL T4 DNA Ligase Buffer, 2.5 uL of NEB Golden Gate Enzyme Mix (BsmBI-v2) Mix (NEB, E1602L), and molbio grade water added to a total of 50uL reaction solution. PCR strip tube with solution was then placed in a pre-warmed thermocycler with a temperature profile of 42C for 60min > 60C for 5min > hold at 4C.

The libraries were then cleaned and concentrated using SPRI using a 1X template:bead ratio (see SPRI cleaning above). Samples were then eluted by adding 15 uL of molbio grade water to the dried beads and keeping ∼12 uL, being sure not to carry through any beads. Next the samples were transformed into electrocompetent cells (Biosearch Technologies, 60242-2). One 0.1cm cuvette (Bio-Rad, 1652089) for each transformation and clean DNA in water were prepared on ice. 1 mL of recovery media (Biosearch Technologies, 80026-1) was transferred to each culture tube in advance. The electrocompetent cells were then thawed on ice. The entire volume of the clean plasmid library (∼10-12 uL) was added to the entire volume of cells (∼50 uL), then mixed gently with a pipette tip 10 times while avoiding bubbles. Entire contents of the DNA+Cell solution was then transferred to the cuvette. The cuvette was then gently tapped to drop cell solution between the metal plates while continuing to avoid bubbles. The solution was then electroporated with the following settings on a Bio-Rad GenePulser Xcell machine (bio-rad, 1652660): 10 uF, 600 Ohms, 1800 Volts. Immediately after the electrical pulse, 1000 uL of Recovery Media was added to the cuvette, then cells were resuspended by pipetting up and down, and the solution was transferred to the 15 mL culture tube (Falcon, 352057). They were recovered at 37 C for 1 hour in an incubator shaking at 300 rpm. Samples were then transferred to 100 mL of LB broth (Corning, 46-050-CM) + Carbenicillin (C2135, Teknova) in 500 mL culture flask (corning, 431401) and incubated at 32.5 C at 160 rpm overnight (16 hrs). Next we plated dilutions to estimate total transformation efficiency as a quick validation. A 0.5 mL aliquot from the culture flask was pipetted into a 1.5 mL eppendorf tube and then diluted twice in recovery media by 1:100 so that the final dilution from the 2 mL recovery solution (before it went into the large 500 mL flask) was 1:10,000 and 1:100,000 respectively. Each dilution was then seeded into carbenicillin agar plates (Teknova, L1011) by adding the full volume of the tube to the plate and then distributing the solution by shaking with seeding beads (Cole-Parmer, UX-01850-33). Plates were then incubated overnight (∼16 hrs) at 37 C. The colonies were counted the next day and coverage was calculated by:

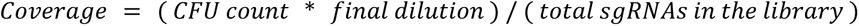

Calculated coverage was compared to expected coverage for any major deviations. The remaining was grown in 100 mL cultures in a 500 mL flask at 30 C overnight at 300 rpm and harvested 16 hours post-transformation. Midiprep was then performed on the samples using NucleoBond Xtra EF plasmid purification, with a final elution volume of 1500 uL. Samples were stored at -20C for long term storage. Dialout and plasmid library distributions were verified by deep sequencing.

### Lentivirus Production

Lenti-X HEK293T cells (632180, Takara) were thawed and cultured in 10% FBS (Gemini, 100-800) in high glucose DMEM (gibco, 10569-010) for at least 1 week before transfection. Cells were passaged every 3 to 4 days when they reached 70-80% confluence. On day 0 of transfection, cells were passaged into 6 well plates (Falcon, 353046) at 1 million cells per well and 2mLs per well of media. On Day 1, cells were transfected using an edited version of the lipofectamine 3000 reagents and protocol (ThermoFisher, L3000075). In short, two solutions were first made. The first with 167 uL per well of OptiMEM (gibco, 51985034) and 14.4 uL of L-3000 reagent. The second solution was made with the following values per well: 167 uL of OptiMEM, 1056 ng of psPAX2 (addgene, plasmid #12260), 704 ng of pMD2.G (addgene, plasmid #12259), 1408 ng of plasmid library, and 12 uL of P3000 reagent. For the second mix it was important to add all of the plasmids first before adding P3000, to prevent the plasmids from crashing out of solution. After mixing the two solutions independently by quickly vortexing, we then added the first solution to the second solution, mixed well and then let incubate for 10 minutes. We then added 333 uL of solution drop wise to each well. On day 2 cells were treated with a final concentration of 2mM viralboost (Alstem, VB100) solution in media by adding dropwise to the wells. On day 3, the virus was harvested by collecting the supernatant and straining it through a 0.45um syringe filter (millipore sigma, SLHV004SL). The virus solution was aliquoted into cryovials and immediately frozen at -80C for later use. Virus was titered on A549s at a wide range of concentrations to determine appropriate volumes to achieve an MOI of approximately 0.25 for screens. Titer was measured via flow by percent BFP positive cells and then the following equation was used to scale the transduction for the screen:

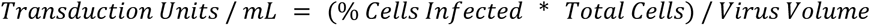

### A549-Cas9 line generation

CAG-Blast-Cas9 plasmid was purchased from Horizon Discovery (Horizon Discovery, CAS10141). A cas9 lentivirus was created as described above. A549s in culture were then transduced with the cas9 lentivirus. Using a selection titer strategy with blasticidin, we selected cells at an MOI of 1. The A549-Cas9 line was then expanded and banked in 10% DMSO (Sigma, D2650-100ML), 10% FBS (Gemini, 100-800), in low glucose DMEM (gibco, 11885-084).

### Cell Culture and Lentivirus Transduction

A549-Cas9 line was thawed into T225 flasks (Thermo Scientific, 159934) and then expanded and selected for Cas9 positive cells with 10 ug/mL blasticidin (gibco, A11139-03) in 10% FBS (Gemini, 100-800) in low glucose DMEM (gibco, 11885-084) for 6 days. They were passaged at 70-80% confluence every 3 or 4 days using TrypLE (gibco, 12604039). On day 6 post thaw, cells were infected with lentivirus plasmid pool at an MOI of 0.3 for 24 hours. Cells were then passaged into multiple T225s for expansion and selected with 1ug/mL puromycin (gibco, A11138-03). Blasticidin selection was discontinued at this point. Transductions and selections were designed to ensure that cell coverage never fell below 300x cells per sgRNA. Cell pellets were collected for next generation sequencing at day 1, 4, 7, and 10 post transduction by spinning down passaged cells at 300g for 5 minutes, aspirating supernatant, and immediately storing at -80C. On day 7 post transduction, cells were seeded into 6-well glass bottom, black plates (Cellvis, P06-1.5H-N) and taken off of puromycin selection. Cells were cultured in 10% FBS in DMEM medium until day 10 post transduction.

### Next Generation Sequencing Sample Processing

NucleoSpin Blood kit (Macherey-Nagel, 740951.50) was used to isolate the genomic DNA. We ran indexing PCR on our NGS samples by creating the following mix per sample: 1x Kapa Hifi readymix HS (Roche, 7958935001), 0.5uM common primer (Table 1), 0.5uM indexing primer (Table 1), 5% DMSO (Sigma, D2650-100ML), 1-10ug of DNA sample, filled up to 100uL with molbio grade water. The thermal profile for the PCR is 95C for 5 min > 95C for 30sec, 56C for 30sec, 72C for 20 sec, 24 cycles > 72C C for 10min > hold at 12C. Samples were split into at least 4 wells of a 96wp PCR plate to minimize jackpotting. DNA was cleaned and quantified using the protocol described in the PCR cleanup section above. Samples were sequenced with the MiSeq V2 with PE50 V2 kit with custom primer according to manufacturer’s recommendations.

Demultiplexing was done by running the BCL files through a BCL-to-fastq pipeline. Fastqs were then used to align sgRNAs to our sgRNA library using an exact match approach of up to 18 base pairs of the 20 total base pairs of the sgRNA. Log2 enrichment was calculated by first getting the cell fraction as:

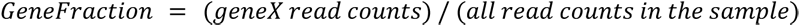

Fold Change was then calculated as:

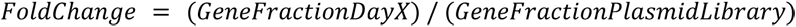

Log2 of Fold Change (FC) was then calculated. Lastly, the FoldChange was then normalized to the intergenic by doing the following:

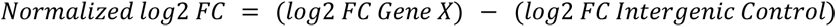

Plots were made using normalized log2 FC.

### MitoProbe

Coordinates for the human mitochondrial ribosome from the Protein DataBase (PDB) (3J9M) were used. We used Visual Molecular Dynamics (VMD) for initial inspection of the PDB. For all other calculations, we used standard python libraries and MDTraj.

Within 3J9M, for all RNA sequences between the lengths of 15 to 25 (inclusive), we computed the SASA for all atoms in the sequence using MDTraj. We then computed the mean, median, and standard deviation of the SASA for all RNA sequences. We then filtered the sequences to only keep sequences that had a fully continuously observed sequence in the PDB. This yielded roughly 4379 sequences though this also contains overlapping sequences (i.e length 15 sequences are in length 16 to 25 length sequences). We sorted these sequences by their mean and median atomic SASA values. We sorted the list by its SASA values and then filtered the sequences by their GC content, absence of repetitive bases, and intra-sequence distances. Out of the final 8 sequences, we picked the first 2 sequences (lengths 20 and 23 base pairs) with the highest SASA (Figure 1) and a large physical separation within the PDB. These selected sequences occupy distinct positions on the mt-ribosome. Oligonucleotides of these sequences were ordered with all uracil (U) bases in the sequence replaced with thymine (T), coupled to 5’ dyes and purified by HPLC (Table 1).

### Cell-Paint POSH protocol

#### Reverse transcription, gap fill, ligation, and amplification

On day 10 post transduction, cells in the 6 well glass bottom plates were fixed by aspirating the medium and replacing it with 4% paraformaldehyde (Electron Microscopy Sciences, 15710-S) in a 1x RNase free PBS solution made from 10x RNase Free PBS (ThermoFisher, AM9625) and molecular biology grade water (Corning, 46-000-CV) for 30 minutes at room temperature. The cells were then washed two times with RNase free 1x PBS.

Plates were permeabilized in 70% ethanol (Sigma Aldrich E7023-500ML) in molbio grade water (Corning, 46-000-CV) for 30 minutes at room temperature. Ethanol was removed by serial dilution with 0.05% Tween in 1x PBS (PBST) by a factor of 1:2 six times to prevent the samples from drying out from ethanol evaporation. Ex: 1mL of ethanol per well, remove 0.5 mL and add 0.5mL PBST - repeat 6x. Samples were then washed fully twice with PBST.

A reverse transcription (RT) solution was then made in molbio grade water (Corning, 46-000-CV) over ice to a final concentration of 1x RevertAid RT Buffer (comes with enzyme), 250uM dNTPs (ThermoFisher, R1122), 0.2 mg/mL BSA (New England Biosciences, B9000S (discontinued)), 1uM Reverse Transcription Primer (see Table 1), 0.8 U/uL Ribolock (ThermoFisher, EO0384), and 4.8 U/uL Revert Aid Reverse Transcriptase (ThermoFisher, EP0452). PBST in all the wells was aspirated and replaced with the RT solution then sealed with sealing film (Applied Biosystems, 4306311) and seal roller (Sigma-Aldrich, R1275-1EA), then incubated overnight (16hrs) at 37 C. Samples were then washed five times with PBST.

Samples were then fixed with a solution of 3.2% paraformaldehyde and 0.1% glutaraldehyde (Sigma-Aldrich, G7651-10ML) in 0.8x PBS for 30 minutes at room temperature. Samples were then washed three times with PBST.

A solution for gap fill, ligation and extension (GF) was made in molbio grade water over ice to a final concentration of 1x Ampligase Buffer (Biosearch Technologies, A1905B), 0.4 U/ uL RNaseH (Enyzmatics, Y9220L), 0.2 mg/mL BSA, 0.1 uM padlock probe (see Table 1), 0.02 U/uL TaqIT (Qiagen Beverly Inc, P7620L), 50 nM dNTPs, and 0.5 U/uL Ampligase (Biosearch Technologies, A3210K). PBST was replaced with the GF solution and the plates were sealed and incubated at 37 C for 5 minutes then 45C for 90 minutes. Plates were then washed twice with PBST at room temperature.

A rolling circle amplification (RCA) solution was made in molbio grade water over ice to a final concentration of 1x phi29 buffer (comes with enzyme), 250 uM dNTPs, 0.2 mg/mL BSA, 5% glycerol (invitrogen, 15514-011), and 1 U/uL phi29 polymerase (ThermoFisher, EP0094). PBST was replaced with the RCA solution, plates were sealed, and incubated at 30°C overnight (16hrs). Samples were then washed twice with PBST. Please note that padlock can only prime to and amplify the sgRNA cassette transcribed with the pol II puromycin and tagBFP transcript, not the pol III transcribed sgRNA. As a result, only the pol II transcript is sequenced by ISS, and not the actual sgRNA, similar to the CROP-seq approach published from Feldman et al^12^.

#### Cell Painting

The clear lid on the Celvis 6 well plates were replaced with a black lid (greinerbio-one, 656199) to protect the samples from light. All staining steps with fluorophores were protected from light as best as possible. If antibodies were used, we first blocked those plates for an hour at room temperature with antibody buffer made of 1% w/v Nuclease free BSA, 0.1% Sodium Azide, 0.1% Nuclease Free TritonX, 1x PBS in molbio grade water. After blocking incubation, blocking solution was exchanged for pS6 primary antibody solution; plates were then sealed and incubated in 4C overnight (∼16hrs). Rabbit anti-human pS6 primary antibody (Cell Signaling Technologies, 2215S) solution was made by diluting pS6 primary Ab 1:250 in antibody buffer solution. Plates were then washed 3 times with PBST (0.05% tween in 1x PBS). All the following staining steps were done on the same day and imaging was started no more than 24 hours after staining for each plate. Any plates not being stained at the time were kept sealed at 4C in PBST until ready to be stained. Staining for all plates was done within 4 days post RCA. Plates were then incubated for an hour at room temperature in secondary antibody mix made of 1:1000 dilution of donkey anti-rabbit DyLight 755 (invitrogen, SA5-10043) in antibody buffer. Plates were then washed three times with PBST at room temperature. Next our mitoprobe mix was prepared, consisting of a final concentration of 0.25uM 12s-1 Cy5 probe (Table 1), 0.25uM 16s-1 Cy5 probe (Table 1), 10% formamide (Fisher Scientific, AC327235000), 10 mM RVC (New England Biosciences, S1402S), and 2x SSC Nuclease free buffer (ThermoFisher, AM9763) in molbio grade water. Cells were then incubated in Mitoprobe mix at 37C for 30 minutes protected from light. They were then washed twice at room temperature with PBST. The CellPaint (CP) staining base was then made in molbio grade water to a final concentration of 1% w/v of Nuclease Free BSA (Sigma-Aldrich, 126609-100GM), 1x HBSS (gibco, 14065-056), and 0.01% Sodium Azide (VWR, BDH7465-2). The CP base buffer was made in bulk (500mLs) and kept at 4C for up to 4 months. A CellPaint mix was made in CP base buffer to a final concentration of 0.33 uM Phalloidin Alexa 568 (ThermoFisher, A12380), 12.5 ug/mL ConA Alexa 488 (ThermoFisher, C11252), 1.5 ug/mL WGA Alexa 555 (ThermoFisher, W32464), 0.5 ug/mL Hoechst (thermofisher, 33342), and 1 U/uL Ribolock (ThermoFisher, EO0384). CellPaint mix was then added to the cells and incubated in the dark at room temperature for 30 minutes. The cells were then washed 5 times with PBST. Cells were immediately imaged or kept in 4C and imaging was started within 24hrs of staining.

Cells were imaged on a Nikon Ti2 Eclipse with 20x 0.45 NA objective. For phenotyping of the 1640 druggable genome screen, the settings and equipment listed below were used (Table 2). The other two screens were done with slightly different filter settings.

**Table 2:**
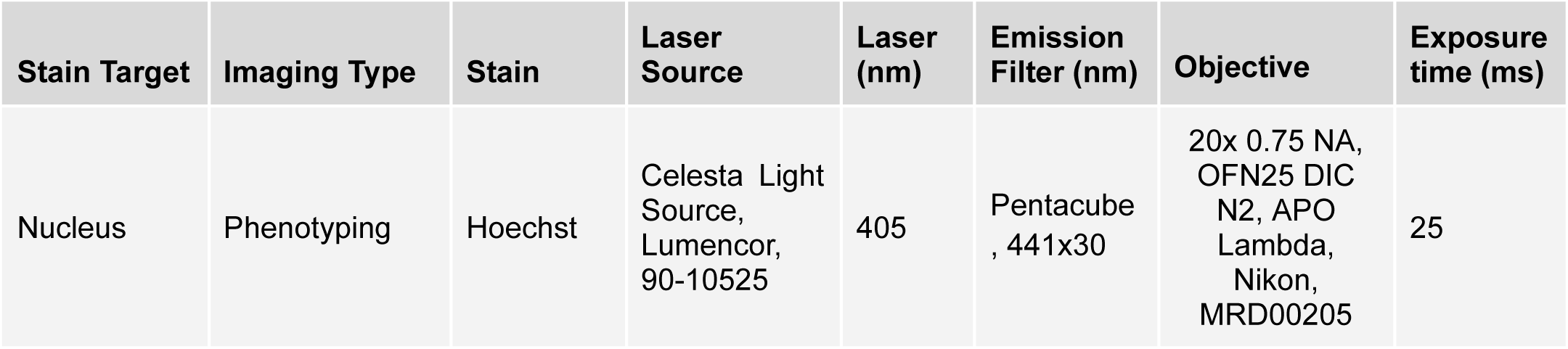
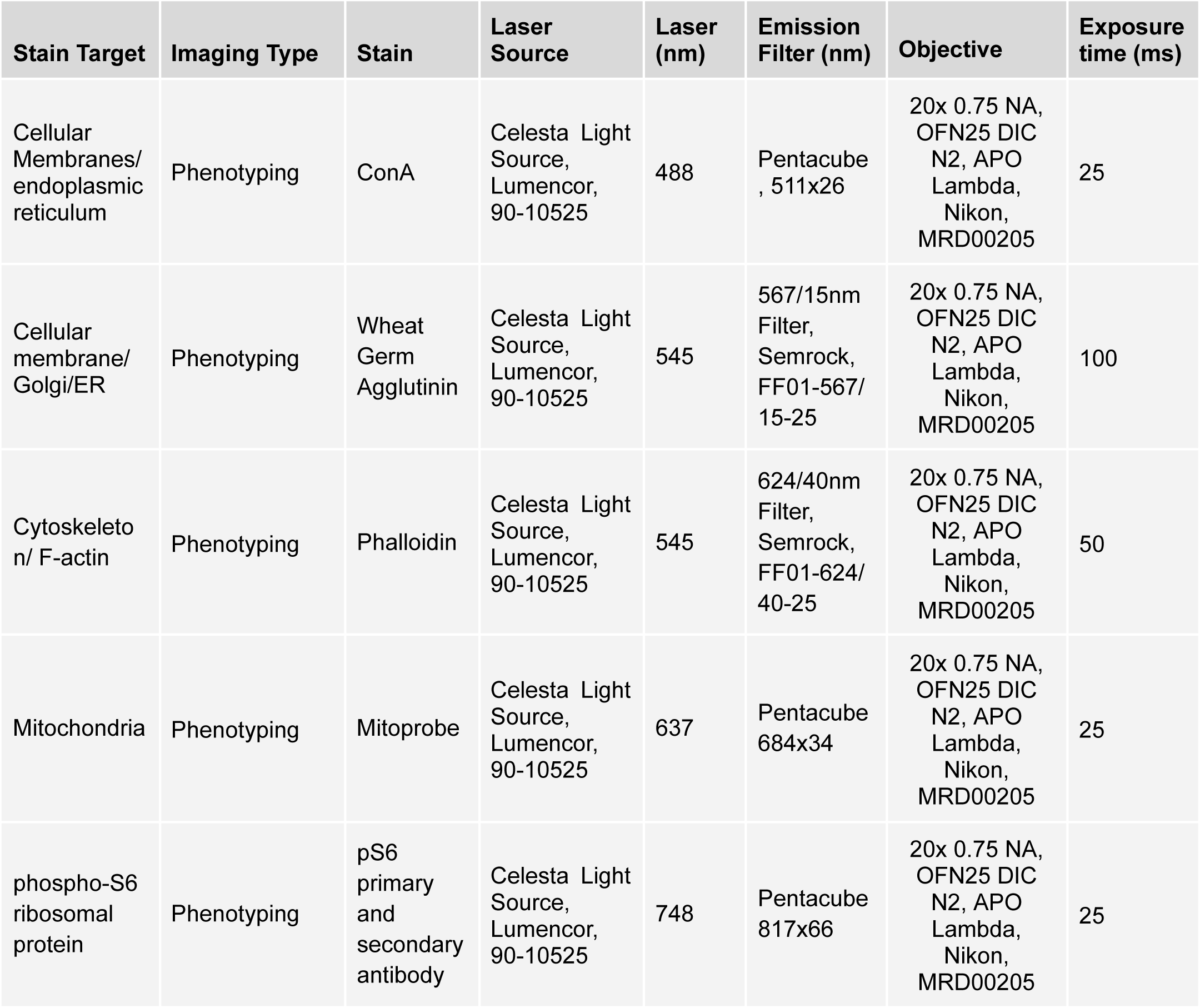
CellPainting + Antibody marker imaging specification.

#### In Situ Sequencing by Synthesis

MiSeq Reagent Kit v2 500-cycles (illumina, MS-102-2003) were used for in situ sequencing by synthesis. Samples were primed for in situ SBS by hybridizing 1uM anchor primer in 2x SSC for 15 minutes at room temperature then washed twice with PR2 incorporation buffer. Incorporation mix for fluorescent labeling of nucleotides was added and incubated for 3 minutes at 60C. The plate was then washed 4 times with PR2 at room temperature, then incubated at 60C in PR2 for 6 mins. Washing and incubation was repeated a total of 2 times. Then the wells were washed twice with PR2. PR2 was then replaced with an imaging buffer made of 200 ng/mL Hoechst (thermofisher, 33342) in 2x SSC and the plate was imaged.

After imaging, cleavage was done to remove fluorescent tags by washing the plate once with PR2, followed by cleavage mix addition and incubation at 60C for 2 minutes. It was then washed 3 times with PR2 at room temperature, then incubated in PR2 at 60C for 2 minutes. It was then washed twice with PR2 at room temperature and incorporation was performed again to add the next base (see above). Steps were repeated until all the desired bases were imaged (about 13 cycles for all screens). Imaging was done using a Nikon Ti2 Eclipse with 10x 0.45NA objective (see Table 3 for more details).

**Table 3:** ISS imaging specification.

| Stain Target | Imaging Type | Stain | Laser Source | Laser Excitation (nm) | Laser Power (%) | Emission Filter | Objective | Exposure time (ms) |
| --- | --- | --- | --- | --- | --- | --- | --- | --- |
| Nucleus | In situ sequencing | Hoechst | Celesta Light Source, Lumencor, 90-10525 | 405 | 15 | Pentacube , 441x30 | 10x 0.45 NA, CFI Plan APO Lambda, Nikon, NRD00045 | 30 |
| Guanine | In situ sequencing | Illumina Incorp. Mix | Celesta Light Source, Lumencor, 90-10525 | 545 | 100 | 567/15nm Filter, Semrock, FF01-567/15-25 | 10x 0.45 NA, CFI Plan APO Lambda, Nikon, NRD00045 | 100 |
| Thymine | In situ sequencing | Illumina Incorp. Mix | Celesta Light Source, Lumencor, 90-10525 | 545 | 100 | 624/40nm Filter, Semrock, FF01-624/40-25 | 10x 0.45 NA, CFI Plan APO Lambda, Nikon, NRD00045 | 100 |
| Adenine | In situ sequencing | Illumina Incorp. Mix | Celesta Light Source, Lumencor, 90-10525 | 637 | 100 | 676/29nm Filter, Semrock, FF01-676/29/25 | 10x 0.45 NA, CFI Plan APO Lambda, Nikon, NRD00045 | 100 |
| Cytosine | In situ sequencing | Illumina Incorp. Mix | Celesta Light Source, Lumencor, 90-10525 | 637 | 100 | 732/68nm Filter, Semrock, FF01-732/68-25 | 10x 0.45 NA, CFI Plan APO Lambda, Nikon, NRD00045 | 100 |

### Druggable Genome Screen Improvements

#### Cell-Paint POSH protocol improvements

To improve the efficiency of our Cell-Paint POSH protocol three modifications were made to improve Cell-Paint and POSH dot quality. First, cell permeabilization was done with 0.1% RNase Free Triton-X (Sigma Aldrich, 93443-500ML) in 1x PBS solution made from 10x RNase Free PBS (ThermoFisher, AM9625) and molecular biology grade water (Corning, 46-000-CV) for 30 minutes at room temperature instead 70% ethanol. This was done to avoid actin degradation by ethanol, which leads to low phalloidin signal. Two washes with PBST were then performed after permeabilization with Triton-X solution. Second, we added a pre-RT fixation step. Briefly, after Triton-X permeabilization, we incubated cells in 1uM reverse transcription primer diluted in PBST for 30 minutes at room temperature. We then replaced the RT primer solution with 3% paraformaldehyde, 0.1% glutaraldehyde in PBST and incubated for 30 minutes at room temperature. We then washed three times with PBST and continued on to reverse transcription as described in the section above. Third, we actively degrade phalloidin signals with ethanol to avoid spectral overlap with the ISS. Briefly, prior to addition of sequencing primer, cells were incubated in 70% ethanol (Sigma Aldrich E7023-500ML) in molbio grade water (Corning, 46-000-CV) for 30 minutes at room temperature. Ethanol was removed by serial dilution with PBST by a factor of 1:2 six times to prevent the samples from drying out from ethanol evaporation. Samples were then washed fully twice with PBST.

#### In Situ Sequencing by Synthesis with Automation

The in situ sequencing process is extremely repetitive, time intensive and tedious, which made it a good candidate for automation (equipment listed in Table 4). In short, the developed system cleaved and incorporated each base pair using a multiflo FX (Biotech, MFXP) for liquid handling, an inheco heater/shaker (inheco, 7100146-A Rev.:04) for heated incubation steps with shaking, a Nikon Ti2 eclipse as described in the previous section for imaging, and a KX2 robot arm (PAA KX-2, KX2-500) to transfer the plate between all three. Briefly, cycle one begins with incorporation of the first nucleotide in Incorporation Mix (Illumina MiSeq), followed by incubation with shaking at 60°C for 3 min, 4 washes with PR2 buffer (PR2, Illumina MiSeq), and 6 minute incubation at 60°C with shaking followed by 4 washes was performed a total of 3 times. The plate is then imaged on the Nikon Ti2. Each subsequent cycle begins with the addition of Cleavage Reagent (Illumina MiSeq) and incubation for 2 min with shaking at 60°C. This is followed by four iterations of four washes (PR2) and incubation for 2 min with shaking at 60°C. The cleaved plate then undergoes the same incorporation process as is found in cycle 1. This is repeated for sufficient cycles to deconvolute POSH barcodes in the library (typically 13 cycles). A time lapse video demonstration is provided in Supp Data 6.

**Table 4:** Automation equipment list.

| Equipment | Vendor | Model # |
| --- | --- | --- |
| Ti2-E Inverted Microscope Main Body | Nikon | MEA54000 |
| Ti2-N-ND-P PFS Motor Nosepiece | Nikon | MEP59394 |
| Ti2-S-SE-E Motorized Stage with Encoder | Nikon | MEC56120 |
| Ti2-CTRE Ti2-E Controller | Nikon | MEF55037 |
| Ti2-F-FLT-E Motorized Epi Fluor Turret | Nikon | MEV51030 |
| Ti2-S-JS Joystick | Nikon | MEF55705 |
| Ti2-LA-BF Fixed Main Branch | Nikon | MEE54860 |
| Ti2-T-BS-S-Eyepiece Tube Base Unit | Nikon | MEB55830 |
| TI2-F-FSS Square Field Stop Slider | Nikon | MEV50030 |
| C-FL Blank Filter Block | Nikon | MXA22030 |
| S-Ti2-Ext External Trigger Cable | Nikon | MXA-22149 |
| S-TI2-DCL Daisy Cable Long | Nikon | MXA22147 |
| USB 2.0 A-B Decvice Cable 15' | Nikon | 97050 |
| Power Cord | Nikon | 79035 |
| CFI Plan APO Lambda 10X | Nikon | MRD00045 |
| CFI Plan APO Lambda 20X | Nikon | MRD00205 |
| TI2-LA-SUE TI2-E Stage Up Kit | Nikon | MED53210 |
| TI2-P0FWBS-E Barrier Filter Wheel STGEUP | Nikon | MEE54410 |
| ORCA-FUSION Gen-III SCMOS Camera C-MT | Nikon (Hamamatsu) | 77054115 |
| Interface Kit F/ORCA Fusion | Nikon | 77054116 |
| C-DA C-MOUNT/ISO Camera Adapter 1X | Nikon | MQD42005 |
| NIS-Elements High Content HC Package | Nikon | MQS34000 |
| Elements: Sutter/.Uniblitz Filterwhl+Shutter | Nikon | MQS41220 |
| NIS-Elements: Wavelength Switcher | Nikon | MQS41930 |
| NIS-Elements: Triggered Device Control | Nikon | MQS42780 |
| NIS-Elements: DAQ TTL/Analog IO | Nikon | MQS41940 |
| NI-BB Plus Analog Triggered Device Hub W/6323 | Nikon | 77023034 |
| Celesta Light Source | Lumencor | 90-10525 |
| 567/15nm Filter | Semrock | FF01-567/15-25 |
| 624/40nm Filter | Semrock | FF01-624/40-25 |
| 676/29nm Filter | Semrock | FF01-676/29/25 |
| 732/68nm Filter | Semrock | FF01-732/68-25 |
| Rapid Filter Switcher | Nikon | MEE54410 |
| Robot Arm | PAA KX-2 | KX2-500 |
| Heater/Shaker | Inheco | 7100146-A Rev.:04 |
| MultiFlo FX Base | BioTek | MFXP2 |
| MultiFlo Secondary Peri | BioTek | 7210010 |
| MultiFlo FX Cassettes | BioTek | 7170011 |
| MultiFlo Washer Module | BioTek | 1260023 |
| MultiFlo 6w Washer Manifolds | BioTek | 1260009 |
| MultiFlo Dual Syringe Dispenser | BioTek | 7210008 |
| SVCE Manifold, 6w 7-degree Assembly | BioTek | 1260537S |
| Overlord Software | Overlord | Version (15.0) |
| MultiFlo Control Software | BioTek | LHC - v5.0 |
| Air Table | Newport | VIS3036-PG2-325A |
| Custom Table | LRG | NA |
| Custom Enclosure | LRG | NA |
| OKO-Lab Drivers<br>(https://www.oko-lab.com/public/okolib/drivers/) | Oko-Lab |  |

### Computational Methods

### Data Capture and Storage

The images acquired on the microscope are exported in raw tiff format and the corresponding metadata for each image as specified in Table 5 are recorded.

**Table 5:** Relevant metadata that are recorded for each acquired image.

| Metadata | Description |
| --- | --- |
| plate_barcode | Barcode of physical plate |
| measurement_id | Identifies an imaging plate run |
| well_row | Row of well in plate |
| well_column | Column of well in plate |
| field_index | Field of view sequential |
| plane_index | Z position sequential. |
| pixel_size_x_um | Size of pixel on detector along the first minor direction |
| pixel_size_y_um | Size of pixel on detector along the first major direction |
| stage_position_x_um | Location of stage as reported by microscope odometry in the first coordinate in microns. |
| stage_position_y_um | Location of stage as reported by microscope odometry in the second coordinate in microns. |

### CellPaint Image Processing

The raw fluorescence field of view images acquired from the microscope have uneven illumination due to camera and vignetting artifacts. In order to create a flat-field illumination image, the raw images are scaled by an illumination correction function as estimated from the mean over field of view images followed by gaussian smoothing as described ^87^. The illumination corrected images are then min-max percentile (min percentile = 0.1, max percentile = 99.9) scaled based on per-channel intensity percentiles computed from plate-level intensity histograms and converted to 8-bit unsigned-integer type (*min value = 0, max value = 255*) for subsequent processing.The field of view images are segmented using CellPose ^88^ to obtain single cell and nuclei boundaries. The field of view images are then cropped around the centroids of each of the segmented nuclei and masked by the corresponding cell segmentation mask to create tiles with a single cell in context.

### *in-situ* sequencing by synthesis Image Processing

The sequencing by synthesis procedure described generates a dataset in which the full plate is imaged several times, with 4-color stationary dots showing variable fluorescent signatures corresponding to A/C/T/G that need to be converted to sequencing base calls.

#### Baseline

The baseline methodology for base calling and sequencing was obtained from Feldman et. al, that first computes locally registered and aligned images across all the SBS cycles followed by blob detection that requires manual fine tuning of parameters.

#### Proposed Methodology

We propose a three-stage ISS methodology that involves first computing base call locations, transforming and projecting the base calls per cycle to the first SBS cycle and stitching of base calls by nearest neighbor matching.

##### (i) Base Calling using convolutional neural network

In order to obtain the base call identities and locations per cycle, we trained a 3-layer fully convolutional network [(*Conv*2*d*, *BatchNorm*2*d*, *ReLU*) *x* 2, *Conv*2*d*, *Sigmoid*] that takes as input the 4-channel SBS image and predicts a 4-channel base identity probability map. To train the model, we generate a proxy labeled dataset using valid base calls generated from the baseline model. The ground truth 4-channel base location/identity images are created by setting a 3×3 pixel region around each baseline detected base call location corresponding to a base to 1 and all other pixels to 0. The convolutional network was trained with supervision on the proxy labeled dataset by optimizing dice loss with a stochastic gradient descent optimizer as described below (*y* = ground truth base label mask, *p̅* = predicted probability mask).

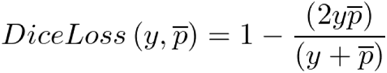

The predicted base call probabilities are converted to predictions by thresholding, labeled by connected components and converted to a table of base call locations and identities per image.

##### (ii) Registration

In order to transform base calls detected in each SBS cycle imaged at different time points to a single acquisition’s coordinate space, we first convert the field of view level base call coordinates (*x_k,f_*, *y_k,f_*) to well coordinate space (*x_k,w_*, *y_k,w_*). We then, compute an odometry transformation matrix (*T_k,1_*) to transform the well coordinates (*x_k,w_*, *y_k,w_*) in each SBS cycle *k* to well coordinates in the first SBS cycle as 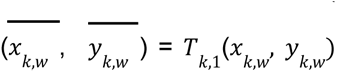. We compute the affine odometry transformation matrix *T* using an FFT based phase cross correlation algorithm ^89^ on a small center crop region of fiduciary markers (Hoechst) between the acquisitions.

##### (iii) Transformation of Base Calls and Barcode Stitching

The base calls computed in each cycle are all transformed to the first SBS cycle using the odometry transformation matrix computed above. The transformed base calls are then chained together based on a KD Tree Nearest Neighbor matching algorithm across cycles to generate a sgRNA barcode readout in the coordinate space of the first SBS cycle.

#### Evaluation and Quality Metrics

The performance and quality of detected base calls and barcodes were assessed using the following metrics

##### (i) Signal to Total Ratio per Cycle

The fidelity of POSH dots throughout several sequencing cycles was measured using the STR equation below for all detected amplicons in each SBS cycle.

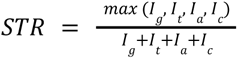

where *I_g_*, *I_t_*, *I_a_*, *I_c_* are the demultiplexed intensities of base fluorophores the detected base locations

##### (ii) Percent of Cells with a Valid Barcode

We define a valid barcode as a stitched barcode that matches with a barcode in our perturbed sgRNA library. For each methodology, we compute the percentage of cells with a valid barcode as the ratio of # of cells with any valid barcode (including multiple valid barcodes) to the total number of segmented cells in the dataset.

### Single Cell sgRNA identity assignment

To assign sgRNA barcode identities to cropped single cell images obtained from CellPaint, we transform the barcode locations, sgRNA sequence and their corresponding gene KO identities from the first cycle of SBS to the CellPaint acquisition space using the same odometry transformation methodology described in the registration section above. The barcodes in CellPaint acquisition space are then assigned to single cell contexts using the cell boundaries and then mapped to corresponding the cropped tiles to obtain a labeled perturbation dataset of single cell images.

### Explicitly Engineered Image Featurization (CellStats)

To localize the signal, we capitalize on the two segmented areas and derive features that are computed in the nucleus, the cytoplasm, and also the perinuclear regions by iteratively dilating the segmentation mask corresponding to the nucleus. First, we start by capturing characteristics of the pixel intensity empirical distributions corresponding to each localization mask. In particular, we extract percentiles from the empirical distributions at discrete steps, along with their means, variances, and standard deviations. Additionally, we compute pairwise Pearson and Spearman correlations among the different channels in our assays. Second, focusing on the localization masks of the nucleus and the cytoplasm, we capture characteristics of each cell’s geometry. Key attributes of cell geometry include the area of the nucleus and the cytoplasm and its convexity ^90^. Lastly, we extract features that have been heavily utilized by the computer vision community to characterize the textures emerging in the nucleus and the cytoplasm from the selected types of staining. In particular, statistics derived from suitable wavelet transforms and region covariance descriptors are employed towards capturing such information. Detailed lists of the features in each of the aforementioned categories are provided in Supp Data 2.

### Self-Supervised Learning of Single-Cell Image Representations

Self-supervised learning pertains to methodologies that are capable of learning latent space representations over input distributions without having an explicit label assigned to them. Recently, self-supervised learning methods [DINO, SimCLR, CLIP] have been shown to improve generalization and quality of learned representations as compared to supervised learning methods ^45,91,92^. In this work, we use DINO ^45^ a self-supervised learning technique that uses knowledge distillation between student 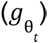 and teacher 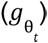 networks parameterized by θ*s* and θ_t_ respectively to learn rich representations from images. Both networks are trained to output a probability vector over *K* dimensions computed as:

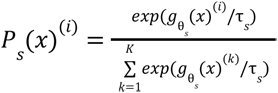

where τ*_s_* is the temperature parameter that controls the sharpness of the output distribution. The student network is trained by minimizing the cross-entropy loss over outputs (as given by Eq below) with the teacher network using stochastic gradient descent.

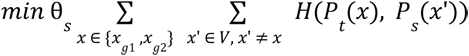

where *H*(*a*, *b*) = − *a log b*, *x_g_*_1_ and *x_g_*_2_ are augmented global views and *V* is a set of augmented local crop views of the input images. The teacher network is updated using an exponential moving average of past weights of the student network as given by:

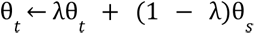

Our motivation to choose DINO as our self-supervised framework stems from their superior performance in k-Nearest Neighbor benchmarks on ImageNet dataset as reported ^45^. We also use a Vision Transformer architecture ^93^ to parameterize both the student and the teacher network parameters.

#### Baseline Pre-trained Vision Transformer

We utilize an ImageNet pre-trained DINO Vision Transformer model (*ViT-small*) with a patch size of 8 trained as our SSL baseline. The pretrained model weights were obtained from [https://github.com/facebookresearch/dino/tree/main]. In order to adapt the pre-trained model trained on 3-channel inputs to 5-channel CellPaint fluorescence image inputs, we re-parameterized the first layer of the model as:

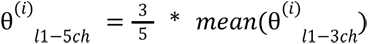

#### Training DINO ViT for Fluorescence Microscopic Images

##### (i) Dataset Preparation

The training dataset was created from the single cell masked and cropped tile images from the 300-gene and 1640-gene CRISPR KO screens for CP-dino-300 and CP-dino-1640 respectively. The dataset contained 1,585,396 cells for CP-dino-300 and 1,559,011 cells for CP-dino-1640. The tile images were then normalized by subtracting by global mean and standard-deviation per channel computed over a small subset of the training images.

##### (ii) Fluorescence Image Augmentation

In order to construct the distorted global and local views of images for DINO training, we modified the set of augmentations to be more relevant for fluorescence microscopic datasets. Some of the key changes are (a) we removed the scale augmentations for both global and local views as all our microscopic images were imaged at a fixed magnification and physical scale. (b) added random rotation augmentations to our images, (c) removed solarization, color jitter and (d) added defocus, coarse dropout and dropout augmentations to generate our global and local views. We also restrict the local crop regions to be within the single cell region to avoid local crops of masked out background regions.

##### (iii) DINO-ViT training

We trained a ViT-small model with patch size = 8, number of global crops = 2, number of local crops = 8 on 4 nodes x 8 NVIDIA-V100 GPUs per node (32 GPUs) for 100 epochs over the 300-gene MoA or the 1640-gene druggable genome (with or without pS6) datasets.

##### (iii) DINO-ViT embedding aggregation

We aggregate gene KO embeddings is as follows: 1) we extract single-cell representations; 2) represent each sgRNA perturbation by the median over all cells of that sgRNA perturbation; 3) represent each gene perturbation as the mean of sgRNA representations corresponding to each gene KO; 4) compute pairwise cosine similarity between gene KO representations.

### Feature Normalization

In order to adjust for intensity variations and batch effects introduced by assay and imaging systems, we normalize each feature embedding by the feature embeddings of *non-targeting* controls in each well. We employ Robust Center-Scale Normalization where for each well, we subtract all the embeddings by the median of *non-targeting* controls in the well and scale by the median absolute deviation of the *non-targeting* controls.

### Feature Aggregation

For extrinsic evaluation of the usefulness of embeddings and understanding biological relationships from embeddings, we first filter the embeddings based on their phenotype score computed by a linear binary classification model between *nontargeting* and every targeting gene KO. We then compute a single sgRNA KO representation by taking the median over all the embeddings corresponding to that sgRNA KO. The gene-KO level aggregate representations are obtained by taking a mean over all the representations of sgRNAs targeting the gene.

### Differential Morphology Analysis and Gene Enrichment Analysis

In several cases, more focused analysis of a single knockout or single feature was necessary. In the case of gene-wise differential morphology analysis, the Z-scored cell populations containing the given knockout, as well as the non-targeting/intergenic control cells within the same pool, were subset from the rest of the pool. Between these two samples, a two-sample Komogorov-Smirnov (K-S) test was applied for each feature with the Z score of the sub-population relative to controls. Due to the large number of features, as well as their redundancy, clear trends in significantly-changed features were annotated; more specific feature results were determined for all genes within the Druggable Genome Screen and created as a report file. A similar approach was applied in the case of feature-wise gene enrichment analysis, in which, for the given feature, each subset of cells receiving the same genetic knockout was compared to non-targeting/intergenic via a two-sample two-tailed K-S test.

### Comparison to FACS Binned 1-Dimensional Analysis

We sought to compare how POSH-based single-cell data on a single feature would compare to a simulated binned FACS screen using the same data. For the single cell resolution based analysis, K-S testing and mean Z were determined as described above for mean cytoplasmic pS6 intensities. To simulate a FACS screen using a pseudo-binning approach, we used the well-normalized Z score matrix and determined the 0-15%, 15-30%, 70-85%, and 85-100% percentile bins, and assigned all cells within the screen to these respective bins. We then assumed perfect sequencing and assignment of the sgRNAs within cells that had been assigned to the bins. MAUDE analysis ^94^ was then conducted to predict the likely distribution shift of cytoplasmic phospho-S6 within each knockout subset, leading to predicted mean Z and FDR for each gene.

### Evaluation of Single-Cell Embeddings by Linear Classifical Model

We evaluate the intrinsic representational power of the different embedding methodologies by measuring their performance on downstream predictive tasks. We evaluate the embeddings by their performance on multiple binary classification tasks for discriminating between non targeting controls and samples corresponding to a gene KO. For each gene KO we train a Elastic Net Logistic Regression model with the task of binary classification between nontargeting and gene KO samples with a leave-one-well-out cross validation schema for all the wells in the experiment. The binary prediction area under the receiver operating characteristic curve (AU-ROC) is computed on the validation samples for each gene in every validation split and the median over AU-ROC is obtained as the scoring function for each embedding. We report the number of gene KOs with an AU-ROC > 0.55 and the mean AU-ROC for the subset of genes on prediction tasks for each embedding methodology as an objective metric for evaluation and comparison.

### Evaluation of Biological Meaningfulness of Aggregate Embeddings

#### Similarity of representations across sgRNAs targeting the same gene

To evaluate the meaningfulness of the aggregate embeddings for each embedding methodology, we computed a pairwise cosine similarity of embeddings [Eq below] for all sgRNAs targeting the same gene.

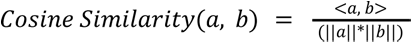

For each embedding method, we then compute the mean sgRNA similarity on a subset of genes that are predictive across all methodologies to rank the embeddings by their meaningfulness in capturing biologically meaningful structure.

#### Evaluating the ability of embeddings to capture known gene-gene relationships in StringDB

While it is difficult to determine “ground truth” in biological datasets, we sought to compare our unbiased screening approach to well-established protein-protein interaction networks in the field. To this end, we used StringDB ^35^, a well-curated network of protein-protein interaction networks based on literature scrubbing, interaction databases, co-expression analysis, and organism transfer. The similarity networks of the genes we screened for were downloaded and used for comparison. For each embedding methodology, using the aggregate gene representations, we compute a pairwise cosine similarity matrix of embeddings. We use the cosine similarity as the strength of similarity/connectivity between two genes learned by the embedding model. We then define the stringDB ground truth connections by thresholding the stringDB *combined score* at different thresholds and compute the area under the receiver operating characteristic curve of the overlap of the embedding cosine similarity matrix with the stringDB network.

### UMAP Visualization of Embeddings

We visualize the aggregate gene-KO embeddings using a UMAP projection with cosine similarity as the metric. The size of each gene-KO node in the UMAP is adjusted based on the similarity of sgRNAs targeting the gene. The gene-KO nodes were colored using Leiden community detection algorithm ^37^ on the UMAP projection.

## Supplementary Data

**Supp Data 1. 124-Gene PoC sgRNA Library**

A detailed summary of the sgRNA library used for the 124-gene PoC screen.

**Supp Data 2. 124-Gene PoC Normalized Cell-Wise Dataset**

The dataset from the 124-gene PoC experiment, including metadata. Values have been normalized (robust center scaled) by non-targeting controls within a given well.

**Supp Data 3. 300-Gene MoA sgRNA Library**

A detailed summary of the sgRNA library used for the druggable genome screen.

**Supp Data 4. 300-Gene MoA screen image dataset (<u>CC-BY-SA license</u>)**

Raw image data of the 300-gene MoA CellPaint-POSH screen

**Supp Data 5. 640-Gene Druggable Genome sgRNA Library**

A detailed summary of the sgRNA library used for the druggable genome screen.

**Supp Data 6. Timelapse video demonstration of automated in-situ sequencing**

## Supplementary Figures

**Figure S1.**
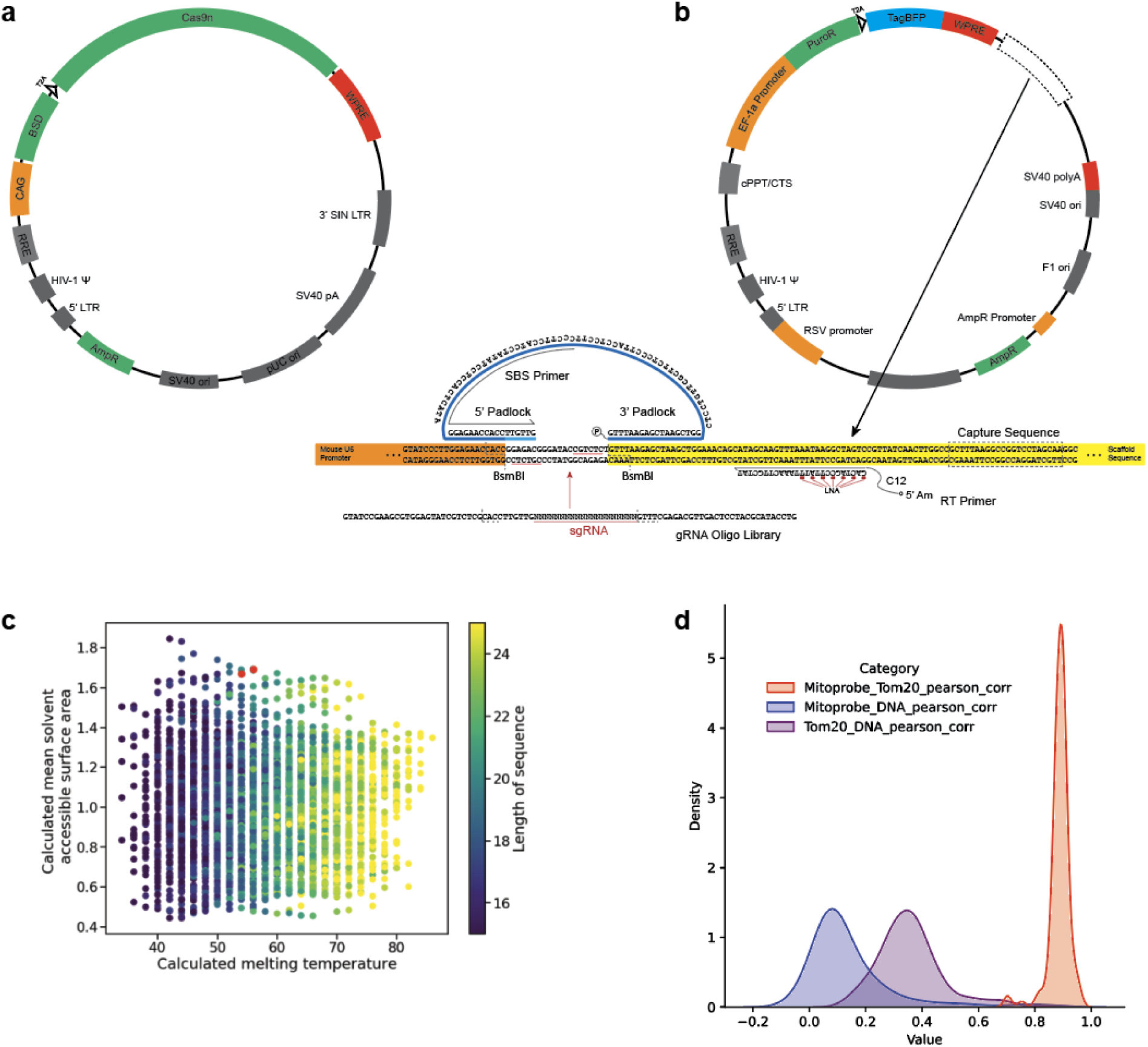
Vectors and supporting data for mitoprobe development. **a.** the vector map of lentivirally-delivered Cas9 nuclease containing blasticidin resistance. **b.** the vector map of the lentivirally-delivered sgRNA. Detailed region shows the mouse U6 promoter (orange), padlock region and sequence (blue), downstream sgRNA scaffold (yellow), and capture sequence (dotted box) for perturb-seq-style compatibility. sgRNA library contains dialout PCR primer flanks and BsmBI restriction enzymes to allow for cloning into the lentiviral backbone. TagBFP is weak and does not interfere with phenotyping or SBS. **c.** optimization of mitoprobe, focusing on a high solvent accessible area and high melting temperature. Selected sequences are highlighted in red. **d.** pearson correlations of Tom20 antibody, Hoechst, and mitoprobe. In designing a mitochondrial stain, a high mitoprobe-Tom20 correlation and low mitoprobe-Hoechst correlation are desired.

**Figure S2.**
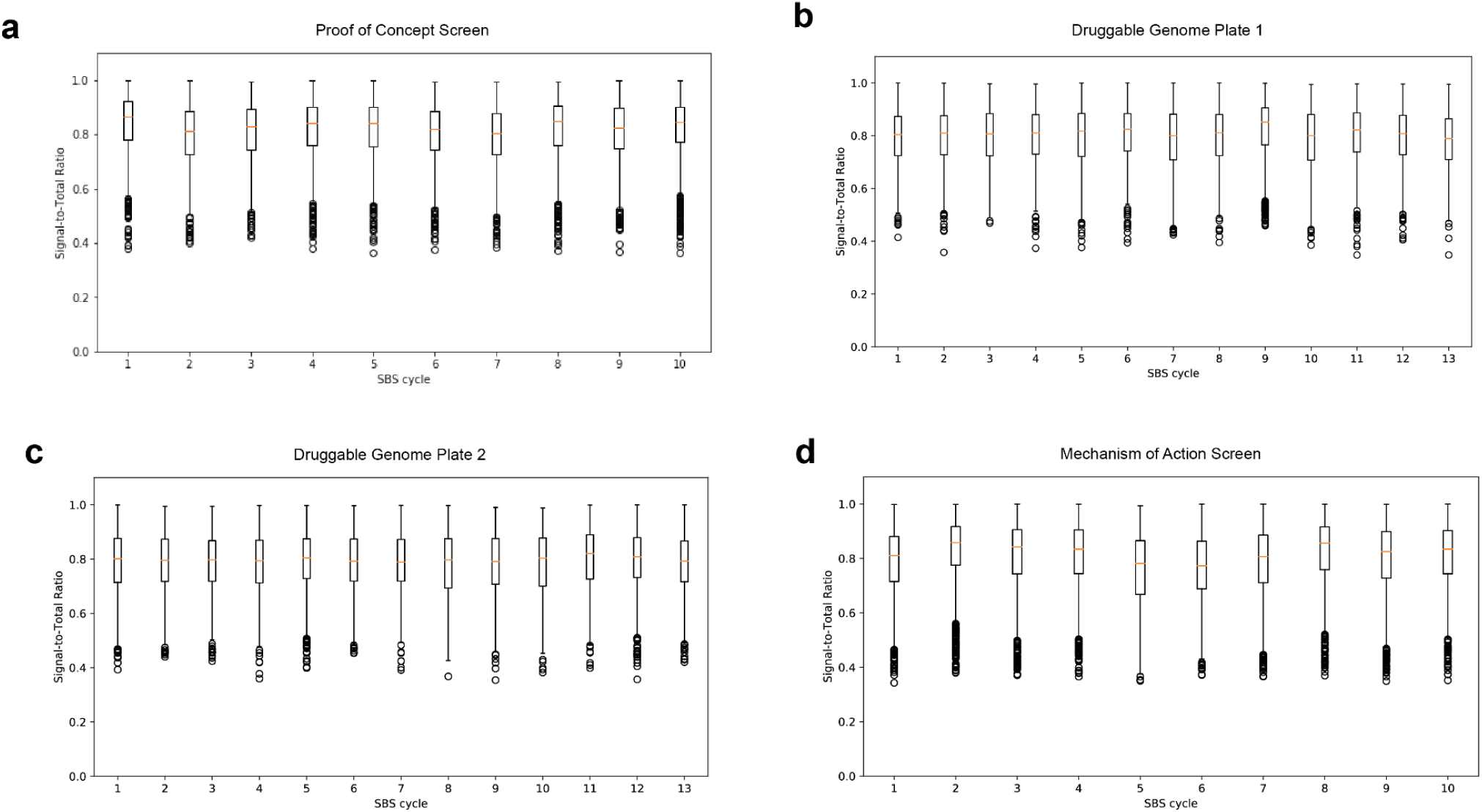
Representative STR plots. **a-d.** Signal-to-total ratio (STR) of the PoC dataset, Druggable Genome dataset, and MoA dataset respectively, showing no significant drop in signal quality across cycles. Poor amplicon generation or sequencing would lead to declining STR due to dimming amplicons, increased background, or phasing of sequences.

**Figure S3.**
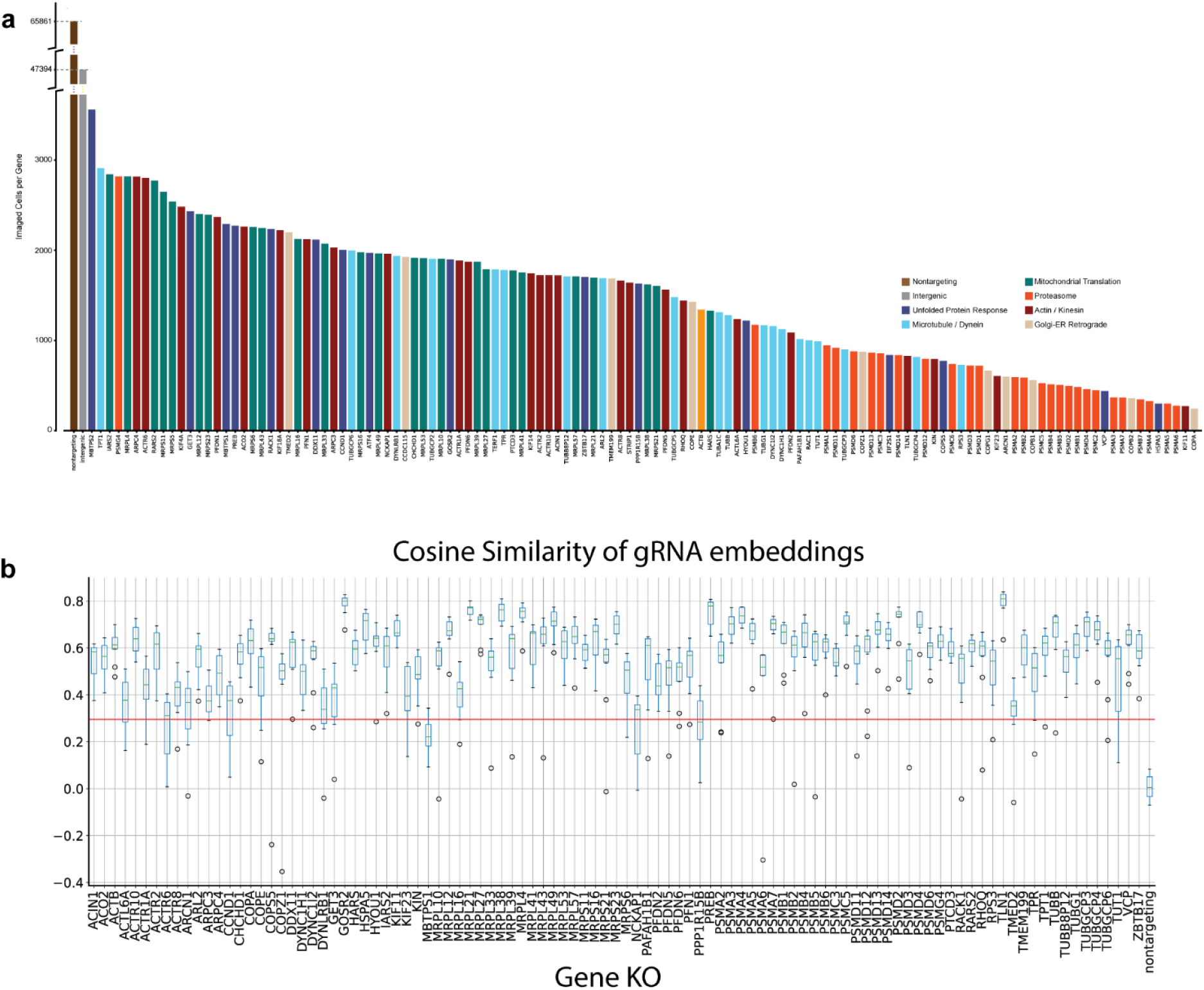
Supporting data from 124-gene proof-of-concept screen. **a.** Total cells detected per gene within the screen. Note the considerable fitness effect created by proteasome inhibition. **b.** sgRNA similarity scores, calculated as the similarities of sgRNAs targeting the same gene. Red line indicates average similarity across all sgRNAs irrespective of targeted genes.

**Figure S4.**
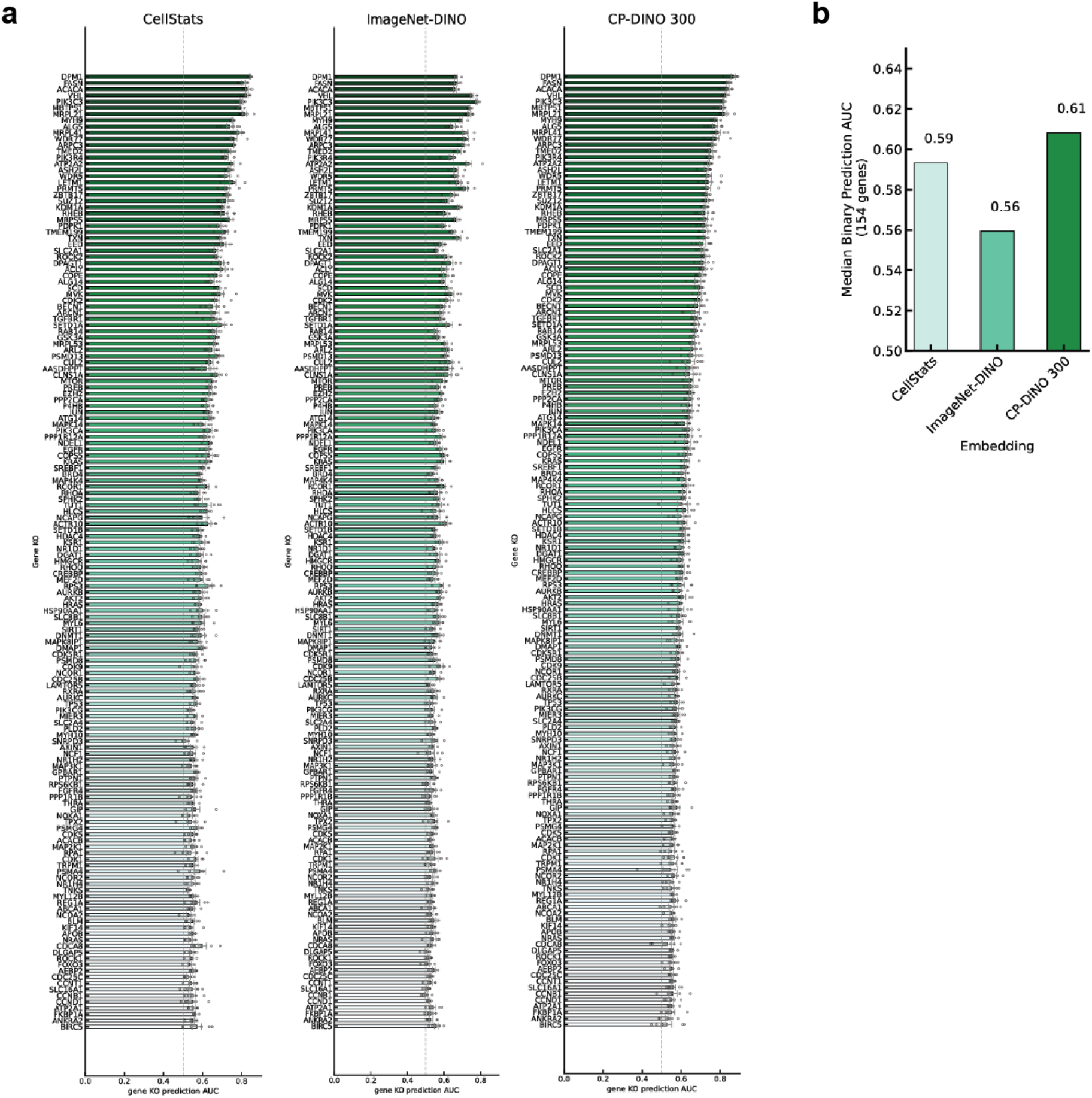
**a**. List of binary classification accuracy (KO vs. WT) for each genetic perturbation ranked by AUC of prediction for all 3 image feature models. **b**. median AUC of prediction from each image feature models.

**Figure S5.**
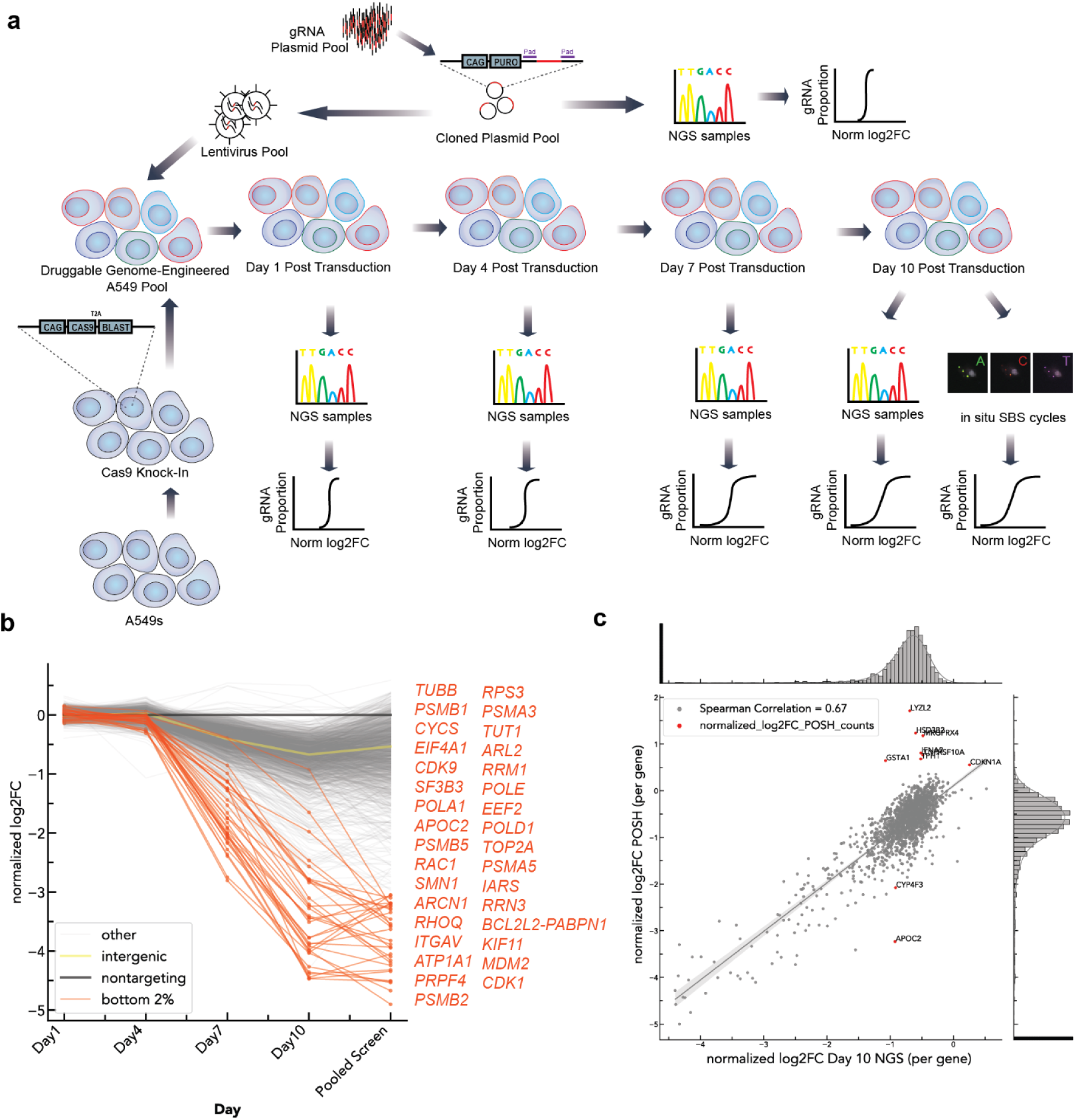
**a**. Workflow diagram of generating time course fitness data for the druggable genome screen. **b**. NGS results for sgRNA distribution (normalized to non-targeting controls), taken through the duration of the screen, along with gRNA distribution calculated by POSH at the final stage of the screen. Orange genes show strong fitness effects that are consistent with NGS validation at time points prior to screening. **c.** scatterplot of the normalized log2fc of gRNA distribution based on the latest NGS sampling and the POSH-identified gRNAs. Genes which show a skewed representation are highlighted.

**Figure S6.**
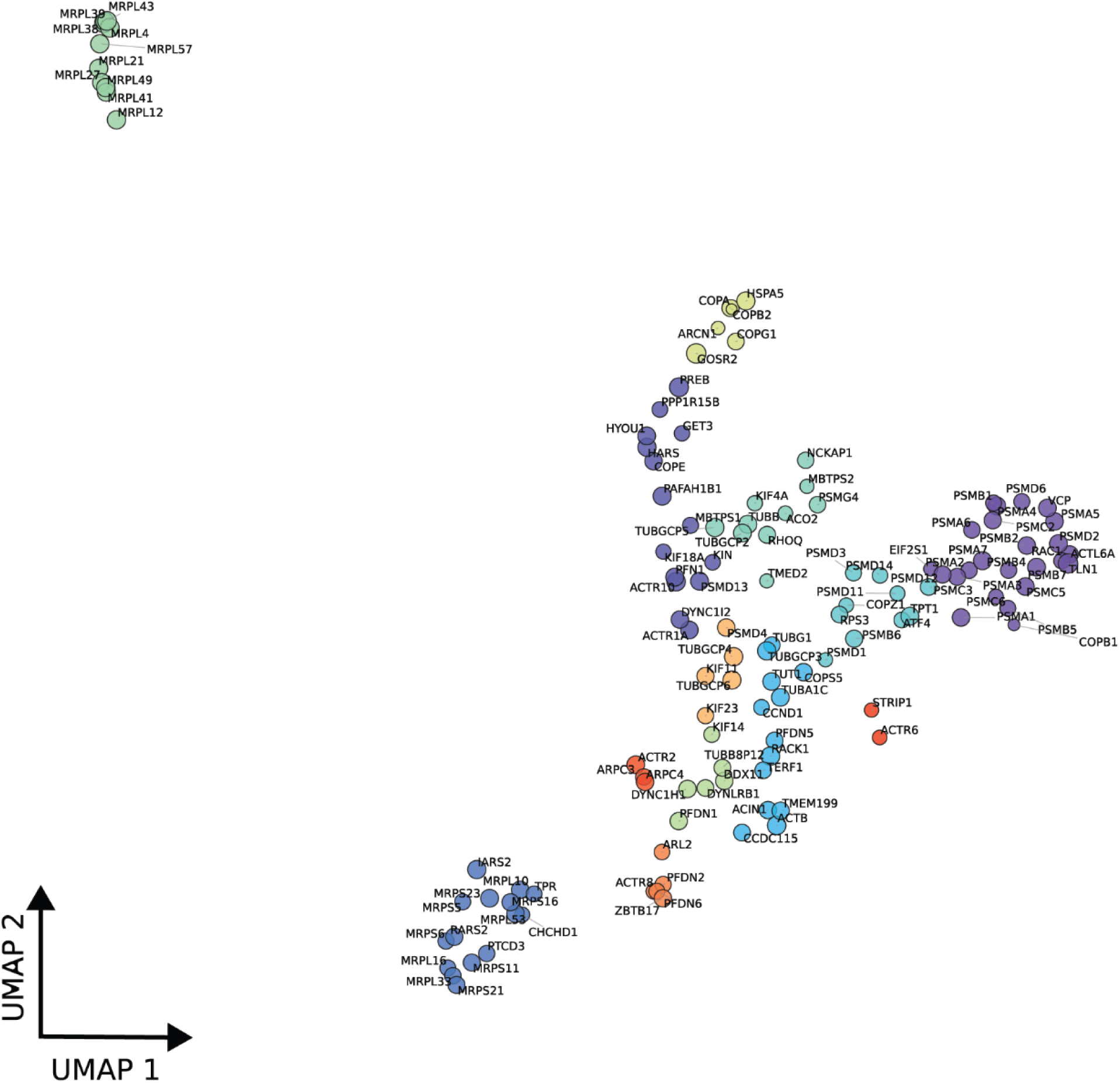
UMAP projection of CP-DINO 1640 features showing genes (filtered by AUC > 0.55) clustered by their biological functions. Colors represent Leiden communities, node size represents the median similarity of aggregate sgRNA embeddings targeting the same gene.

**Figure S7.**
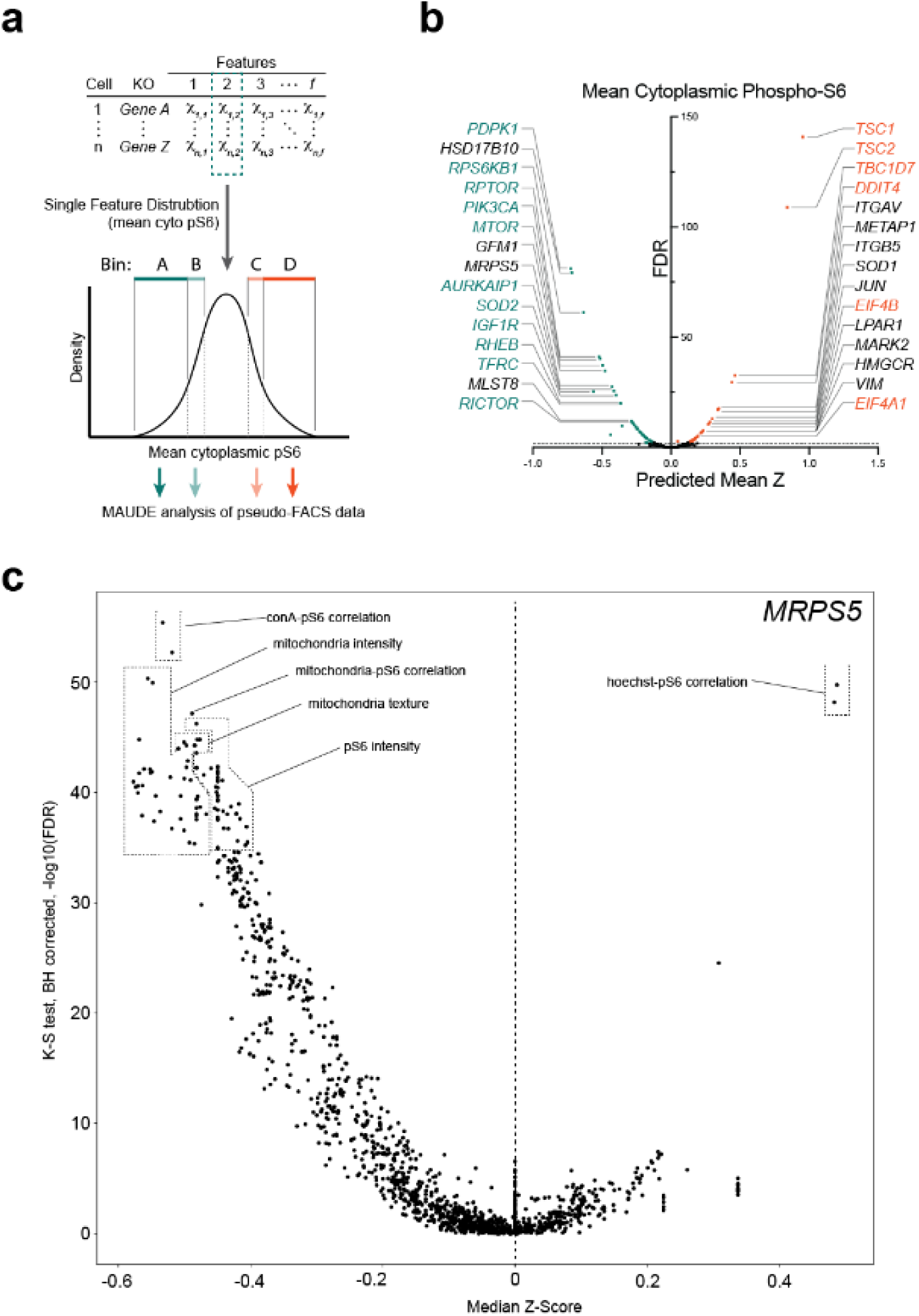
**a-b.** the pipeline and results for simulated pseudo binned FACS data, using MAUDE to predict the likely shift in mean cytoplasmic phospho-S6 caused by each gene’s regulation. While some shifting in order occurred, this analysis resulted in similar top-ranked genes to when the data is analyzed using single cell information. **c.** differential feature analysis on *MRPS5*.

